# A Gαq/11-IP_3_R1-ANO1 pacemaking axis underlies pressure-induced lymphatic chronotropy

**DOI:** 10.1101/2025.10.07.681001

**Authors:** Michael J. Davis, Hae Jin Kim, Min Li, Jorge A. Castorena-Gonzalez, Mary E. Schulz, Grace Pea, Soumiya Pal, Timothy L. Domeier, Joshua P. Scallan, Scott Earley, Scott D. Zawieja

**Author notes:** Correspondence: Michael J. Davis, PhD Department of Medical Pharmacology & Physiology University of Missouri School of Medicine One Hospital Drive, MA415 Medical Sciences Building Columbia, MO 65212.

## Abstract

The frequency of spontaneous lymphatic contractions is exquisitely sensitive to changes in intraluminal pressure, enabling lymph transport to be matched to the filling state of the lymphatic capillary network. The mechanistic basis of this mechanosensitive response was investigated using ex vivo contraction assays of mouse popliteal collecting vessels. Based on electrophysiological and transcriptional similarities between lymphatic muscle cells (LMCs) and arterial smooth muscle, we hypothesized that pressure-dependent chronotropy shares a parallel signaling process with pressure-induced arterial depolarization/constriction. Thus, we probed two potential mechanisms: 1) pressure-induced activation of mechanosensitive cation channels and 2) mechano-activation of GNAQ/GNA11-coupled G-protein receptors (GPCRs) that would generate second messengers to activate those channels. Contraction assays were combined with scRNAseq analysis of the respective targets and use of transgenic mice. Our findings rule out significant roles for the mechanosensitive channels implicated in arterial myogenic constriction as well as channels implicated in ionic pacemaking of other tissues. In contrast, smooth-muscle specific knock out or inhibition of ANO1, IP_3_R1 or GNAQ/GNA11 significantly abrogated pressure-induced chronotropy. These results support a scheme whereby mechano-activation of one or more GNAQ/GNA11-coupled GPCRs generates IP_3_, inducing sarcoplasmic reticulum Ca^2+^ release through IP_3_R1, which drives depolarization through the activation of ANO1 Cl^-^ channels. The major GPCRs expressed in LMCs were identified and ranked by scRNAseq analysis but individual inhibition or knock out the top 7 candidates failed to significantly impair pressure-induced chronotropy. Our results implicate one or more GNAQ/GNA11-coupled GPCRs in mediating this mechanosensitive process, but the specific GPCRs remain to be identified.

## Introduction

The lymphatic system is critically important for the maintenance of interstitial fluid balance. Because the Starling forces governing fluid movement across blood capillaries favor net filtration in most tissues (Levick & Michel, 2010), filtered fluid and protein will accumulate in the interstitium unless it is removed by the lymphatic system. Lymphatic capillaries are optimized to reabsorb that fluid, which can exceed 8 L·day^-1^ (Renkin, 1986). Once reabsorbed, lymph is transported by a combination of passive and active transport mechanisms through the collecting lymphatic vessel network, first to lymph nodes and eventually back into the venous system. Active lymph pumping relies on the spontaneous contractions of collecting lymphatic vessels—events that are initiated by action potentials (APs) in lymphatic muscle cells (LMCs). These contractions are relatively large in amplitude, with ejection fractions often exceeding 80% and sufficient to propel lymph through a system of one-way valves that ensure its unidirectional movement (Davis *et al*., 2025). Active pumping accounts for the majority of lymphatic transport in the lower legs of humans during quiet standing (Olszewski & Engeset, 1980) and facilitates lymph return against adverse hydrostatic pressure gradients that develop under gravitational loads (Davis *et al*., 2012; Scallan *et al*., 2016; Li *et al*., 2022). An important feature of the active lymph pump system is the exquisite sensitivity of the spontaneous contraction frequency of collecting lymphatic vessels to changes in their intraluminal pressure, enabling lymph transport to be matched to the filling state of the lymphatic capillary network (Scallan *et al*., 2012). This homeostatic mechanism is termed “pressure-induced lymphatic chronotropy” (Zawieja *et al*., 2019). In popliteal lymphatic collecting vessels of the mouse hindlimb, the spontaneous contraction frequency can increase from 2 to 20 contractions·min^-1^ as pressure is elevated from 0.5 to 10 cmH_2_O, i.e., a 10-fold increase over the physiological pressure range in those vessels (Davis *et al*., 2020; Davis *et al*., 2023a; Davis *et al*., 2023b). Contraction amplitude increases modestly over a portion of the same pressure range (Scallan *et al*., 2012), but modulation of contraction frequency is the primary method for acute adjustments in active lymph transport.

Pressure-induced chronotropy reflects an underlying mechanotransduction process that ultimately involves control of an intrinsic pacemaker in LMCs by intraluminal pressure. Although the specific force sensing mechanism is unknown, contractile, electrophysiological and transcriptional similarities between LMCs and vascular smooth muscle cells (VSMCs) support the hypothesis that pressure-dependent chronotropy shares a parallel signaling process to the myogenic response of VSMCs. The myogenic response refers to the rapid increase in arterial pressure induced by distention of a small artery / arteriole, which then constricts over time until a new, stable diameter smaller than the original diameter is reached (Davis, 2012). A similar constriction often occurs in collecting lymphatic vessels coincident with pressure-induced chronotropy (Davis *et al*., 2009). Pressure-induced arterial constriction is preceded by VSMC depolarization (Harder, 1984) and several lines of evidence support a critical role for the activation of (putative) mechanosensitive cation channels in that process—in particular the involvement of three TRP channel family members: TRPC6, TRPM4 and PKD1/2.

The first evidence for a molecularly identified ion channel involved in myogenic constriction came from the demonstration that single-channel currents in VSMCs isolated from rat cerebral arteries were activated by patch pipette suction and those currents, along with pressure-induced cerebral artery constrictions and depolarizations, were attenuated after downregulation of TRPC6 with antisense oligonucleotides (Welsh *et al*., 2002). TRPM4 channels were likewise implicated after inward, TRPM4-like currents in rat pial artery myocytes were found to be activated by membrane stretch, and because antisense- or siRNA-mediated knock down of TRPM4 significantly attenuated pressure-induced depolarization and myogenic tone development in those arteries (Earley *et al*., 2004b; Gonzales *et al*., 2014). A recent study has confirmed that SM-specific TRPM4 knockout significantly impairs myogenic tone (Zhu *et al*., 2026). In contrast, results from *Trpc6*^-/-^ and *Trpm4^-/-^* mice tended to contradict the above studies in finding no differences or even *enhanced* arterial myogenic constriction in the global knock out animals (Dietrich *et al*., 2005; Masathar *et al*., 2010; Schleifenbaum *et al*., 2014). Evidence also supports roles for PKD2 (previously described as TRPP1) channels in myogenic constriction, but that, too, is controversial. Jaggar and colleagues reported that shRNA-mediated down-regulation of *Pkd2* attenuated pressure-induced depolarization and myogenic tone in rat pial arteries (Narayanan *et al*., 2013) and that pressure-induced constriction was blunted in hindlimb arteries isolated from Myh11-CreER^T2^;*Pkd2^f/f^* (smooth muscle knockout, smKO) mice (Bulley *et al*., 2018). However, a study by Honoré and coworkers suggested that stretch-activated cation currents and myogenic constriction in mouse mesenteric artery smooth muscle were mediated by PKD1 channels (previously termed TRPP2), with PKD2 (previously termed TRPP1) channels playing a counteracting role (Sharif-Naeini *et al*., 2009). Subsequent studies showed that myogenic tone is impaired in *Pkd2* smKO mice (Bulley *et al*., 2018) but not in *Pkd1* smKO mice (Bernardelli *et al*., 2026).

Recent work points to G-protein-coupled receptors (GPCRs) rather than TRP or other ion channels as the primary mechanosensing elements in the arterial myogenic response. Studies of cardiomyocyctes first demonstrated that the angiotensin receptor, AT_1_R, a GNAQ/GNA11-coupled GPCR, could be activated by mechanical stress through a mechanism independent of its ligand (Zou *et al*., 2004). Gudermann and coworkers provided compelling support for this mechanism using patch-clamped HEK293 cells co-expressing AT_1_R and TRPC6 channels, with hypoosmotic swelling used as a mechanical stimulus (Mederos y Schnitzler *et al*., 2008). Two critical findings were 1) that TRPC6 current was activated by hypoosmotic swelling *only* if the cells co-expressed *Agtr1* (Mederos y Schnitzler *et al*., 2008) and 2) that the swelling-induced currents were largely prevented by inverse AT_1_R agonists. *Agtr1* could be replaced by another GNAQ/GNA11-coupled GPCR, or *Trpc6* by another DAG-sensitive TRP channel (Mederos y Schnitzler *et al*., 2008). Subsequent studies from multiple laboratories, using selective pharmacological inhibitors of AT_1_R, acute *Agtr1* knockdown, and/or mice deficient in *Agtr1* or other GNAQ/GNA11-coupled GPCRs, confirmed the basic principle in VSMCs: that pressure-induced constriction of many different types of arteries is transduced primarily, or exclusively, by GNAQ/GNA11-coupled GPCRs (Blodow *et al*., 2014; Harraz *et al*., 2014; Li *et al*., 2014b; Schleifenbaum *et al*., 2014; Pires *et al*., 2017; Bjorling *et al*., 2018; Cui *et al*., 2022). Helix 8 in the GPCR C-terminus was subsequently identified as the essential structural motif endowing mechanosensitivity (Erdogmus *et al*., 2019). The collective data support a model in which phospholipase C is activated downstream from GNAQ/GNA11-coupled GPCRs in response to elevated pressure, leading to increased production of DAG and IP_3_/Ca^2+^ release, which then activate TRPC6 and TRPM4 channels, respectively, to depolarize the cell and increase the open probability of L-type voltage-gated Ca^2+^ channels, thereby promoting global Ca^2+^ influx and contraction (Davis *et al*., 2023d). In a similar manner, TRPM4 and PKD2 channels might be activated by Ca^2+^ influx/release downstream from mechano-activation of a GPCR or another Ca^2+^-permeable ion channel such as PIEZO1 (Peyronnet *et al*., 2013) or TRPV4 (Swain & Liddle, 2021).

Whether similar signaling pathways are operative in LMCs is unknown. Each lymphatic contraction is preceded by an approximately linear depolarization from the “resting” potential to the threshold potential, termed diastolic depolarization, which is the primary determinant of AP / contraction frequency (Zawieja *et al*., 2025). Thus, lymphatic pressure elevation does not produce a stepwise increase in baseline LMC membrane potential (von der Weid *et al*., 2014; Davis & Zawieja, 2018) but rather an increase in the diastolic depolarization slope and a reduction in the time required to reach the threshold for AP firing (Zawieja *et al*., 2018b; Zawieja *et al*., 2019). Previously, we found that regulation of the diastolic depolarization slope was mediated largely by activation of the Ca^2+^ activated chloride channel ANOCTAMIN 1 (ANO1, also referred to as TMEM16A). ANO1 is not thought to be intrinsically mechanosensitive, but regulated by Ca^2+^ and, in LMCs specifically, by IP_3_ receptor 1-mediated Ca^2+^ release from sarcoplasmic reticulum (SR) (Zawieja *et al*., 2019; Zawieja *et al*., 2023), possibly downstream from one or more mechanosensitive GPCRs. However, other mechanisms may also be involved because *Ano1* deletion from smooth muscle or pharmacological inhibition of ANO1 does not completely eliminate spontaneous APs or contractions (Zawieja *et al*., 2019). The primary goal of the present study was to test the roles of putative VSMC mechanosensitive ion channels, other channels implicated in the regulation of pacemaking in different cell types, and GNAQ/GNA11-coupled GPCRs in pressure-induced chronotropy of popliteal lymphatic vessels. An advantage in studying this particular aspect of lymphatic vessel mechanotransduction is the high sensitivity of the spontaneous contraction frequency to pressure, increasing ∼10-fold over only a 5 cmH_2_O pressure range in popliteal lymphatics compared to relatively modest levels of myogenic constriction (50% change over a 60 mmHg pressure range) even in the most myogenically reactive arteries (Davis *et al*., 2023d). As part of our strategy, we sought to make maximum use of transgenic mice to avoid non-specific effects of pharmacological inhibitors, particularly those used to block TRP channels.

## Results

### Quantification of pressure-induced chronotropy

Our previous studies of the mechanism underlying pressure-induced chronotropy used mouse IALVs (Zawieja *et al*., 2019; Zawieja *et al*., 2023). However, the contraction frequency of IALVs changes only ∼1.7 fold over the pressure range 0.5 to 10 cmH_2_O, compared to the much larger dynamic range (≥ 10-fold) characteristic of popliteal lymphatics. Thus, for this study we used the responses of popliteal lymphatics to assess pressure-induced chronotropy, in anticipation of being able to detect even very subtle changes in that response after deletion of specific genes. An example of pressure-induced chronotropy in a WT mouse popliteal lymphatic is illustrated by the original recording in **Fig. 1A**. The basal contraction frequency was 11 min^-1^ at the equilibration pressure of 3 cmH_2_O, with frequency slowing to 3.2, 2.7 and 0.5 min^-1^ when pressure was lowered to 2, 1 and 0.5 cmH_2_O, respectively.

**Figure 1.**
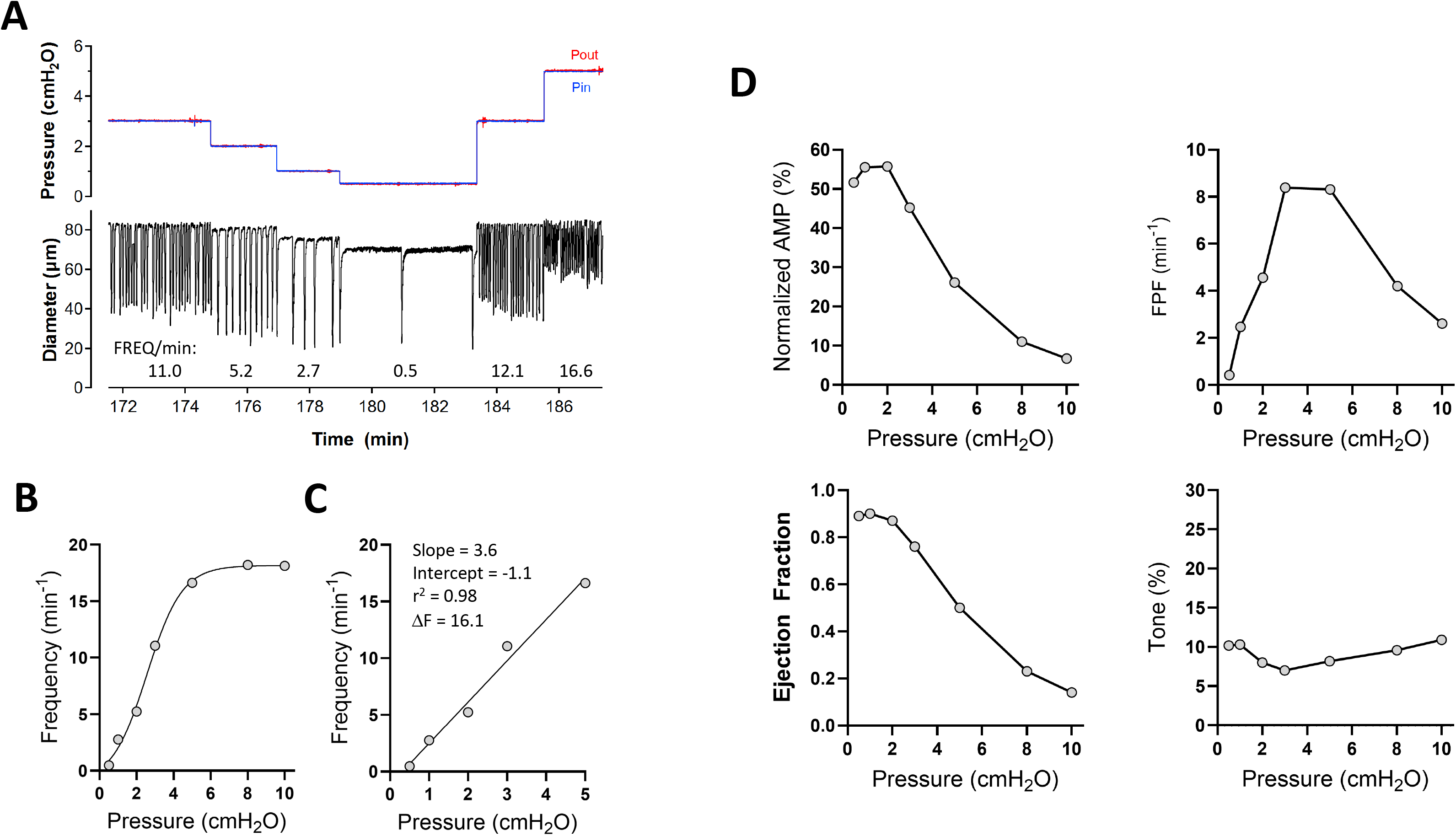
Quantification of the F-P relationship. Procedure used for quantitative assessment of pressure-induced chronotropy. **A**) Representative recording of pressure and diameter changes in a WT popliteal lymphatic vessel during simultaneous inflow (Pin) and outflow (Pout) pressure steps between 0.5 and 5 cmH_2_O. Frequencies stated at the bottom of the trace were determined off-line using a custom peak detection program written in LabVIEW. **B**) Plot of frequency vs pressure for the vessel in A, showing how frequency reaches a plateau above 5 cmH_2_O. **C**) Fit of the frequency data with a first-order equation and calculation of the frequency difference (ΔF) between 5 and 0.5 cmH_2_O. **D**) Summary of contraction data for this representative WT vessel over the stated pressure range; see Methods for description of parameter calculations.

Frequency rose to 12.1 min^-1^ when pressure was stepped back to 3 cmH_2_O and increased further to 16.6 min^-1^ when pressure was subsequently raised to 5 cmH_2_O. A plot of frequency vs pressure (F-P) is shown in **Fig. 1B** for the entire pressure range (including 8 and 10 cmH_2_O). The data between 0.5 and 5 cmH_2_O were fit with a linear regression, with a slope of 3.6 and a change in frequency of 16.1 min^-1^ between 0.5 and 5 cmH_2_O, as shown in **Fig. 1C**. Other major contraction parameters—amplitude, ejection fraction, estimated fractional pump flow and tone—are shown in **Fig. 1D** for each pressure level.

### Role of ANO1 in pressure-induced chronotropy

Previously, we reported that ANO1 inhibition or SM-specific *Ano1* deletion caused a nearly complete abrogation of pressure-induced chronotropy in mouse IALVs (Zawieja *et al*., 2019). A similar analysis is presented for popliteal lymphatic collectors in **Fig. 2**. scRNAseq analysis confirmed moderate expression levels of *Ano1* (relative to *Acta2*) in >75% of LMCs and to a lesser extent in AdvCs, but not in LECs (**Fig. 2A**). IALVs were used for the scRNAseq analysis because of the larger number of cells available from those vessels and similarities in the mechanisms of pressure-induced chronotropy between the two vessel types are discussed later. Cells associated with collecting lymphatic vessels segregated into 19 distinct clusters/subclusters, including lymphatic endothelial cells (LECs), LMCs, adventitial cells (AdvCs) and immune cells, as previously described (Zawieja *et al*., 2025). Two small clusters reflected contamination by mammary gland ducts, which are present within the tissue in the vicinity of the IALVs. AdvCs contained subclusters of adventitial fibroblasts with some expressing multiple cell markers of pluripotency (Zawieja *et al*., 2025).

**Figure 2.**
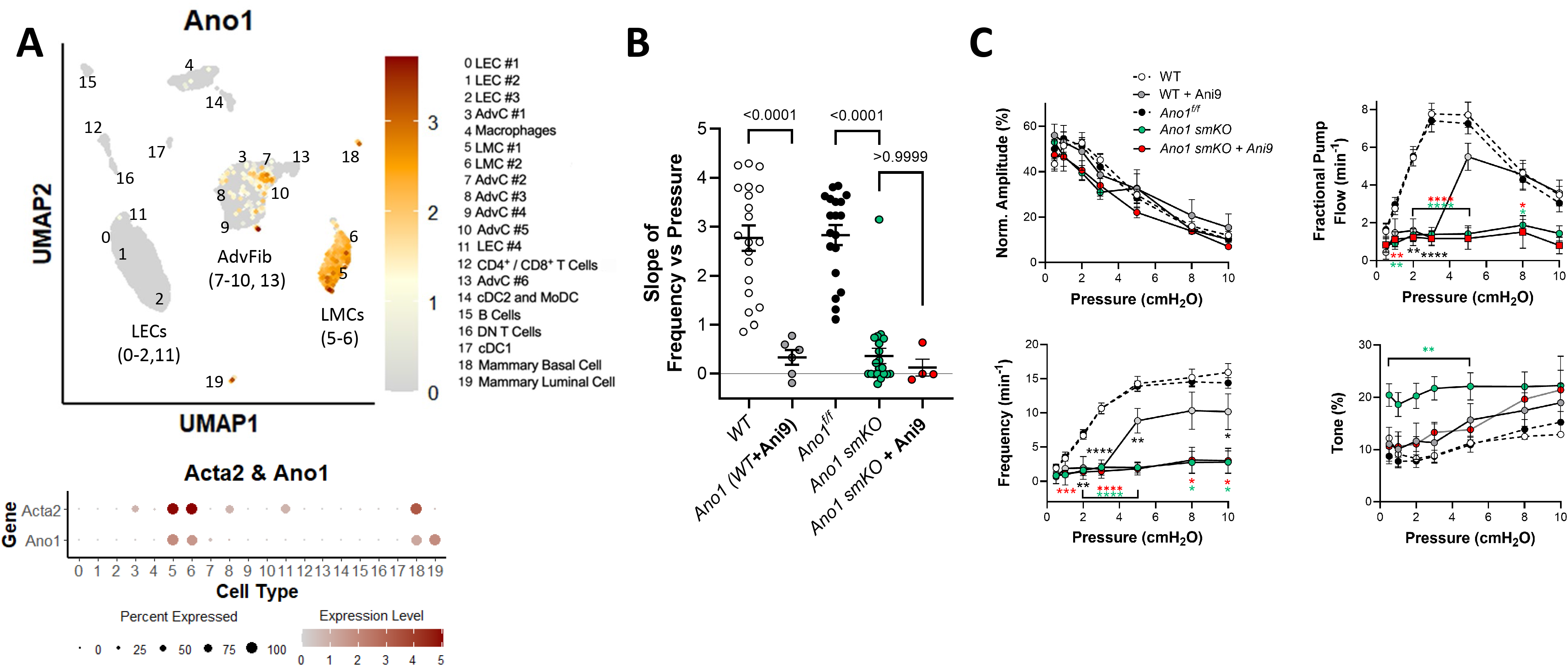
Consequences of ANO1 deletion and/or inhibition. **A**) UMAP plot of *Ano1* expression in the various cell populations within the walls of murine IALVs that were thoroughly cleaned prior to dissociation and scRNAseq analysis, as described in (Zawieja *et al*., 2025). The LMC cluster, composed of at least two subclusters of 978 cells, was identified by the expression of canonical smooth muscle cell markers *Acta2* (SM α-actin), *Myh11* (myosin heavy chain 11) and *Itga8* (integrin alpha 8). Gray color in this and subsequent figures represents *Acta2* expression, red/brown colors in this panel indicate *Ano1* expression. Bubble plot shows Ano1 expression in LMCs and other cell clusters in terms of percent cells in an individual cluster expressing *Ano1* (size of dot) and the *Ano1* expression level relative to *Acta2* (color of dot). Abbreviated names of the clusters are listed on the right. **B**) Slope of frequency vs pressure (F-P) for WT (C57Bl/6) vessels with / without treatment by the ANO1 inhibitor Ani9 (1 μM), for *Ano1^f/f^* control vessels, for *Ano1* smKO vessels and for the latter treated with Ani9. **C**) Summary plots of contraction parameters as a function of pressure for the 5 respective groups of vessels. Horizontal bars indicate ranges of pressures with same specified statistical significance. WT N=15, n=20; WT+Ani9 N=5, n=6; *Ano1* smKO N=14, n=23; *Ano1^f/f^* N=8, n=16; *Ano1* smKO+Ani9 N=4, n=4.

The F-P analysis for 20 WT popliteal lymphatic vessels (open black symbols) is shown in **Fig. 2**. C57Bl/6 mice served as the WT control group for many comparisons because all transgenic strains used in the present study (with one exception) were derived from, or backcrossed into, that background. The average slope (± SEM) of the F-P relationship between 0.5 and 5 cmH_2_O was 2.8 ± 0.3 for WT vessels (**Fig. 2B**), equating to an average frequency difference of 12.1 ± 1.0 contractions·min^-1^ over that pressure range. The average r^2^ value for the linear regressions was 0.93 ± 0.02, indicating a consistently high quality of the curve fits and confirming that the slope was approximately linear between 0.5 and 5 cmH_2_O. *Ano1^f/f^*vessels served as another control group (filled black symbols), and these showed a nearly identical average slope for their F-P relationship (2.9 ± 0.2). The role of ANO1 in pressure-induced chronotropy of mouse popliteal lymphatic vessels was then tested after pharmacologic inhibition of ANO1 with Ani9, one of the most selective small molecule inhibitors of ANO1 (Hwang *et al*., 2016). Initial assessment of the Ani9 concentration-response relationship (from 30 nM to 10 μM) in WT popliteal vessels revealed that 3 μM Ani9, the concentration previously used to inhibit ANO1 in IALVs (Zawieja *et al*., 2019; Zawieja *et al*., 2023), did indeed reduce the basal contraction frequency of popliteal lymphatics (in the lower pressure range), but also caused a loss of tone and marked reduction in their contraction amplitude (not shown)— suggestive of possible off-target effects; therefore, for subsequent F-P protocols we used a lower concentration (1 μM) of Ani9, which had insignificant effects on contraction amplitude and tone (**Fig. 2C**) but significantly reduced the average slope of the F-P relationship from 2.8 ± 0.3 to 0.3 ± 0.2 (p<0.001; **Fig. 2B**).

We then tested the effects of SM-specific *Ano1* deletion by comparing the F-P relationship of popliteal lymphatic vessels from tamoxifen-treated Myh11-CreER^T2^*;Ano1^f/f^* (*Ano1* smKO) mice to *Ano1^f/f^* control mice. The results are also shown in **Fig. 2B**. The average slope of the F-P relationship between 0.5 and 5 cmH_2_O for *Ano1* smKO vessels was 0.4 ± 0.2, significantly lower (p<0.001) than that for *Ano1^f/f^* vessels. Complete sets of the other contraction parameters for untreated and Ani9-treated WT popliteal lymphatics, Myh11-CreER^T2^*;Ano1^f/f^*and *Ano1^f/f^* popliteal vessels, are shown in **Fig. 2C**, including the average frequencies at each pressure in the lower left plot. The frequency of Ani9-treated vessels was significantly reduced at most pressures but increased at higher pressures. Normalized contraction amplitudes were unchanged by Ani9. Tone was slightly but not significantly higher in Ani9-treated vessels. After *Ano1* deletion, frequency was reduced to <3 contractions·min^-1^ at all pressures, in contrast to Ani9-treated WT vessels in which frequency was comparably reduced at low pressures but not at elevated pressures (**Fig. 2C**). *Ano1* smKO vessels had significantly elevated levels of tone at 5 of 7 pressures, suggesting that Ani9 treatment may have interfered with tone development in a way that *Ano1* deletion did not. Collectively, these results show that deletion / inhibition of ANO1 significantly impaired pressure-induced chronotropy in popliteal lymphatic vessels; however, SM-specific *Ano1* deletion failed to completely abolish spontaneous contractions in 16 of 21 *Ano1* smKO vessels. To test the possibility that residual pacemaking activity in *Ano1* smKO vessels might reflect incomplete recombination by the inducible Myh11-CreER^T2^ driver, *Ano1* mRNA levels in IALVs were quantified using qPCR analysis. Smooth muscle cells were digested from IALVs of Myh11-CreER^T2^*;Rosa26^LSL-mTmG^* or Myh11-CreER^T2^*;Rosa26^LSL-^ _mTmG_;Ano1^f/f^*mice and purified by FACS using the GFP signal. *Ano1* message (referenced to α-actin levels) in LMCs from Myh11-CreER^T2^*;Rosa26^LSL-mTmG^;Ano1^f/f^* vessels was undetectable compared to that in LMCs from Myh11-CreER^T2^*;Rosa26^LSL-mTmG^* vessels (0 ± 0 and 1006 ± 80, respectively; mean ± SD, n=3). As a further test for possible residual ANO1 activity in *Ano1* smKO vessels, we treated another group of *Ano1* smKO vessels with Ani9 (1 μM), and those results are shown in **Fig. 2B-C**. Pacemaking activity persisted even after combined deletion and inhibition of ANO1, without significant further reductions in frequency at any pressure, suggesting that residual pacemaking in *Ano1* smKO vessels was *not* due to incomplete *Ano1* knock down. Examples of the contraction patterns observed in *Ano1^f/f^* and *Ano1* smKO vessels are shown in **Figure 3**; in the KO vessel, the basal contraction frequency was low (2.8 at P=3 cmH_2_O) compared to the control vessel and did not increase further (2.2 min^-1^) at P=5 cmH_2_O. Collectively, these results point to ANO1 as the major component of pressure-induced lymphatic chronotropy but also to the involvement of one or more additional mechanisms, possibly involving other LMC ion channels.

**Figure 3.**
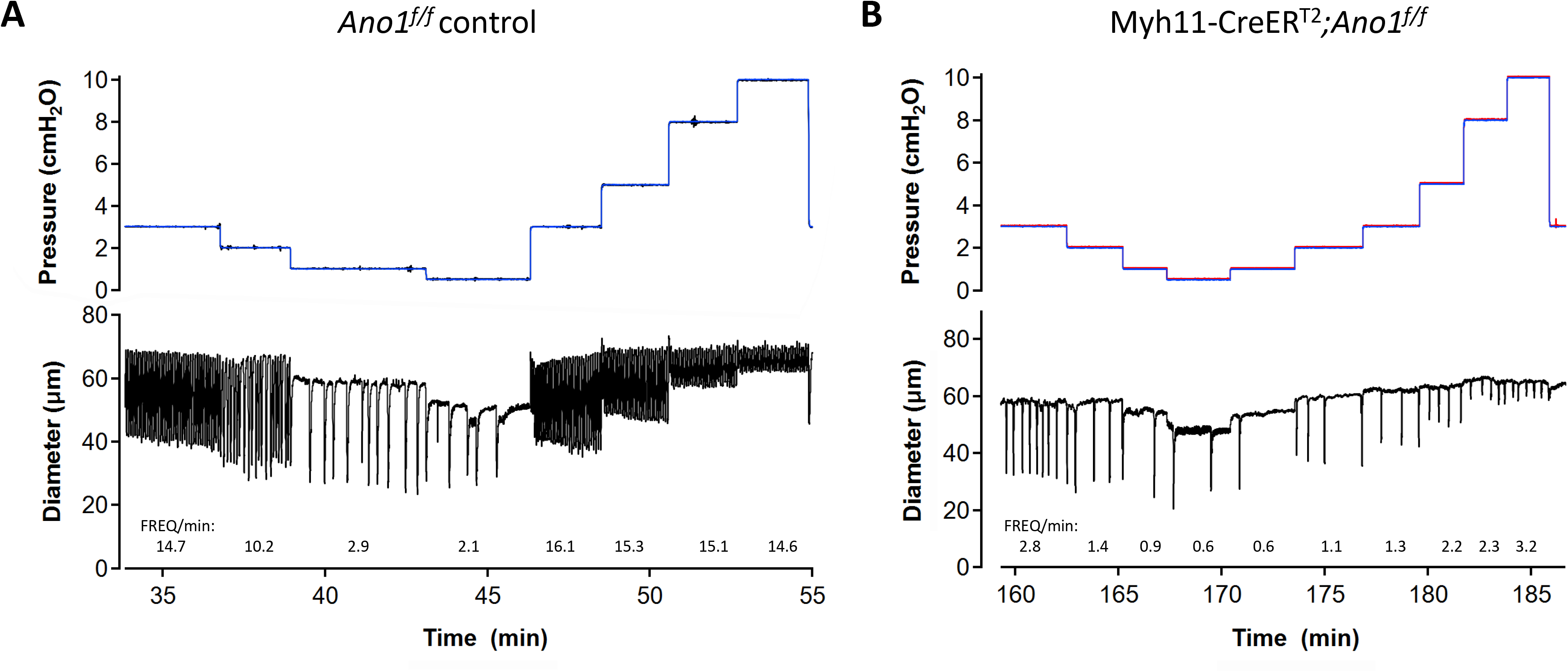
Contraction patterns of Ano1^f/f^ and *Ano1* smKO vessels. A) Spontaneous contractions of an *Ano1^f/f^* (control) popliteal lymphatic vessel at various pressures. Average frequencies at each pressure are listed below the diameter traces. Frequency ranged from 2.1 min^-1^ at 0.5 cmH_2_O to 15.3 min^-1^ at 5 cmH_2_O. B) Spontaneous contractions of a Myh11-CreER^T2^;*Ano1^f/f^* (*Ano1* smKO) popliteal vessel at various pressures. In this particular example, two additional pressure steps between 0.5 and 3 cmH_2_O were imposed (this example represents an early experiment before the protocol was finalized). Frequency ranged from 0.6 min^-1^ at 0.5 cmH_2_O to 2.2 min^-1^ at 5 cmH_2_O.

### Role of PIEZO1

PIEZO1 and PIEZO2 are two ion channels that satisfy the full set of criteria for true mechanosensitivity (Coste *et al*., 2010; Davis *et al*., 2023d). PIEZO2 is expressed primarily in neurons (Woo *et al*., 2014; Woo *et al*., 2015) whereas PIEZO1 is more widely expressed (Murthy *et al*., 2017). PIEZO1 is critically important for shear-stress induced Ca^2+^ signaling in endothelium (Li *et al*., 2014a), required for lymphatic valve and vessel development (Nonomura *et al*., 2018; Choi *et al*., 2019; Choi *et al*., 2024) and for hypertension-dependent arterial wall remodeling (Retailleau *et al*., 2015). The role of PIEZO1 in the arterial myogenic response has not been systematically tested, except for a single report showing that SM-specific *Piezo1* deletion did not alter myogenic tone development in either caudal or rostral cerebellar arteries of mice [Suppl. Figs. 2-3 in (Retailleau *et al*., 2015)]. The possible involvement of PIEZO1 in lymphatic muscle contractility has not been reported. Our scRNAseq analysis revealed strong expression of *Piezo1* message in LECs but very weak expression in LMCs (**Fig. 4A**). *Piezo2* expression was barely detectable in either cell type. Popliteal lymphatic vessels from Myh11-CreER^T2^*;Piezo1^f/f^* (*Piezo1* smKO) and *Piezo1^f/f^*control mice were subjected to the protocol for assessing pressure-induced chronotropy. However, the data revealed no significant difference in the F-P slope between *Piezo1* smKO vessels and *Piezo1^f/f^* control vessels (**Fig. 4B**), suggesting that PIEZO1 channels in LMCs are not required for pressure-induced lymphatic chronotropy. As a further test, we used Nestin-Cre to generate Nestin-Cre*;Piezo1^f/f^ (Piezo1* nestinKO) mice with constitutive deletion of *Piezo1* both in smooth muscle and other cells expressing intermediate filaments. An analysis of vessel responses from *Piezo1* nestinKO mice indicated that there was no significant impairment in pressure-induced chronotropy (**Fig. 4B**). Although we did not suspect the endothelium would be involved in pressure-induced chronotropy, we also generated and tested Prox1-CreER^T2^*;Piezo1^f/f^ (Piezo1* lecKO) mice. As expected, there was no significant difference in the average F-P slope of vessels from *Piezo1* lecKO mice compared to tamoxifen-treated *Piezo1^f/f^* controls (**Fig. 4B**). Complete sets of contraction data for popliteal vessels from *Piezo1* smKO are shown in **Fig. 4C** and revealed no significant differences in any of the lymphatic contractile parameters at any pressure compared to their floxed controls.

**Figure 4.**
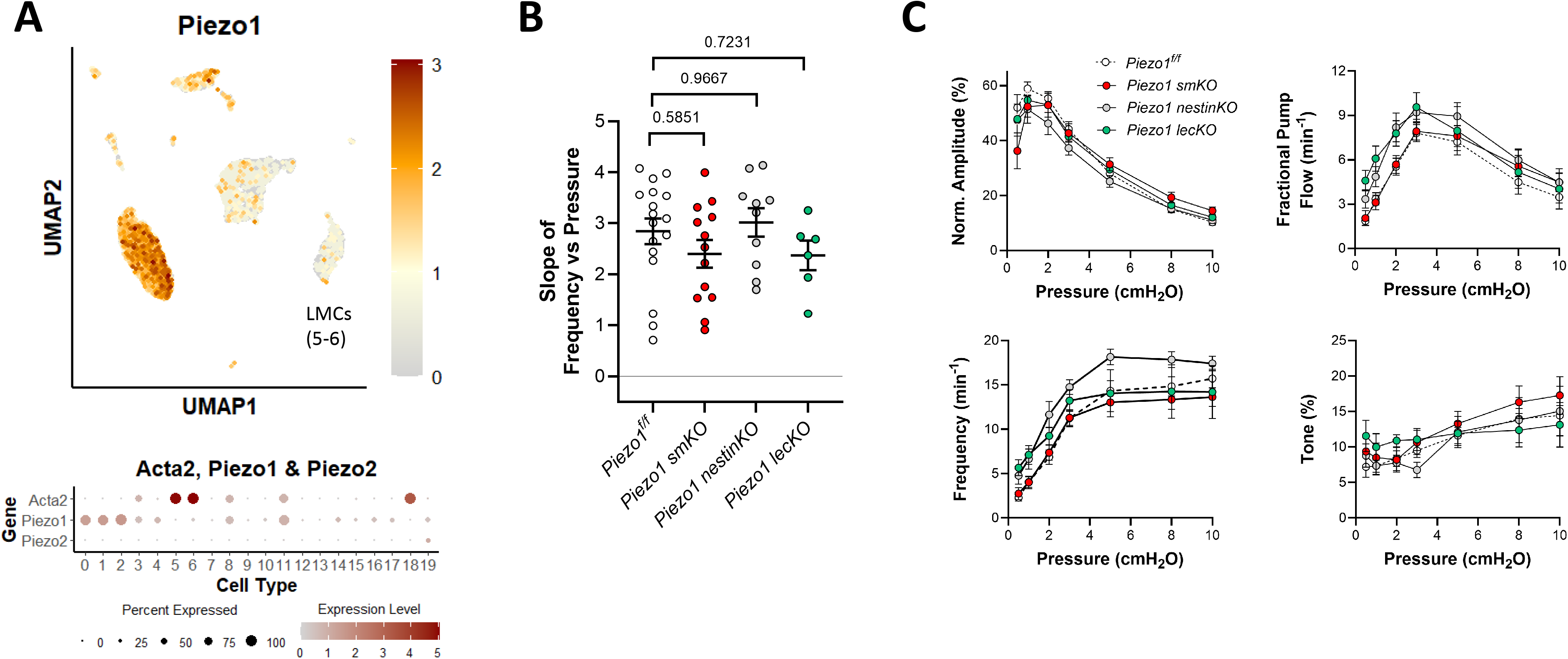
Consequences of *Piezo1* deletion. **A**) UMAP and bubble plots of *Piezo1 and Piezo2* expression in the various IALV cell clusters. **B**) Slope of F-P relationship for popliteal lymphatics from *Piezo1^f/f^, Piezo1* smKO, *Piezo1 nestin KO*, *Piezo1* lecKO mice. **C**) Summary plots of contraction parameters as a function of pressure for the 4 respective groups of vessels. *Piezo1^f/f^* N=9, n=17; *Piezo1* smKO N=7, n=13; *Piezo1 nestin KO* N=5, n=10; *Piezo1* lecKO N=5, n=6.

### Roles of TRPC6 and TRPC3

TRPC6 is thought to be a mechanosensitive cation channel whose activation underlies pressure-induced constriction in cerebral arteries (Welsh *et al*., 2002). However, studies of other arteries from *Trpc6^-/-^*mice found either no differences from control vessels or even *enhanced* myogenic responsiveness (Dietrich *et al*., 2005; Schleifenbaum *et al*., 2014). Because global TRPC6 deletion results in compensatory upregulation of TRPC3 (Dietrich *et al*., 2005), a related TRPC family member with similar properties that is also activated by DAG, it is possible that TRPC3 could compensate for loss of TRPC6. The expression patterns for *Trpc6* and *Trpc3*, based on scRNAseq analyses, are shown in **Fig. 5A**. *Trpc6* was expressed at modest levels in ∼50% of LMCs and was almost exclusively confined to LMCs. *Trpc3* was expressed at very low levels in only a few LMCs (from control mice). We tested the effects of global TRPC6 deletion (*Trpc6^-/-^*), global TRPC3 deletion (*Trpc3^-/-^*) and global deletion of both isoforms (*Trpc6^-/-^;Trpc3^-/-^*) on pressure-induced lymphatic chronotropy. As these mice were all generated in the SV129 background, control measurements were performed on popliteal vessels from SV129 mice, which showed a *lower* average F-P slope than WT (C57Bl/6) popliteal vessels. There were no significant differences in F-P slope between SV129 controls and *Trpc6^-/-^* or *Trpc3^-/-^* vessels, but the F-P slope of *Trpc6^-/-^;Trpc3^-/-^*vessels was significantly *higher* than that of the SV129 controls (**Fig. 5B**). Complete sets of contraction data for *Trpc6^-/-^;Trpc3^-/-^*vessels, compared to their *SV129* controls, are shown in **Fig. 5C**. *Trpc6^-/-^;Trpc3^-/-^*double knock-out vessels had significantly impaired contraction amplitudes at several pressures and *increased* frequencies and *increased* levels of tone at most pressures. However, relevant to the main objective of this study, there were no significant *impairments* in pressure-induced chronotropy in vessels deficient in *Trpc6*, *Trpc3* or both *Trpc6* and *Trpc3* (**Fig. 5B-C**).

**Figure 5.**
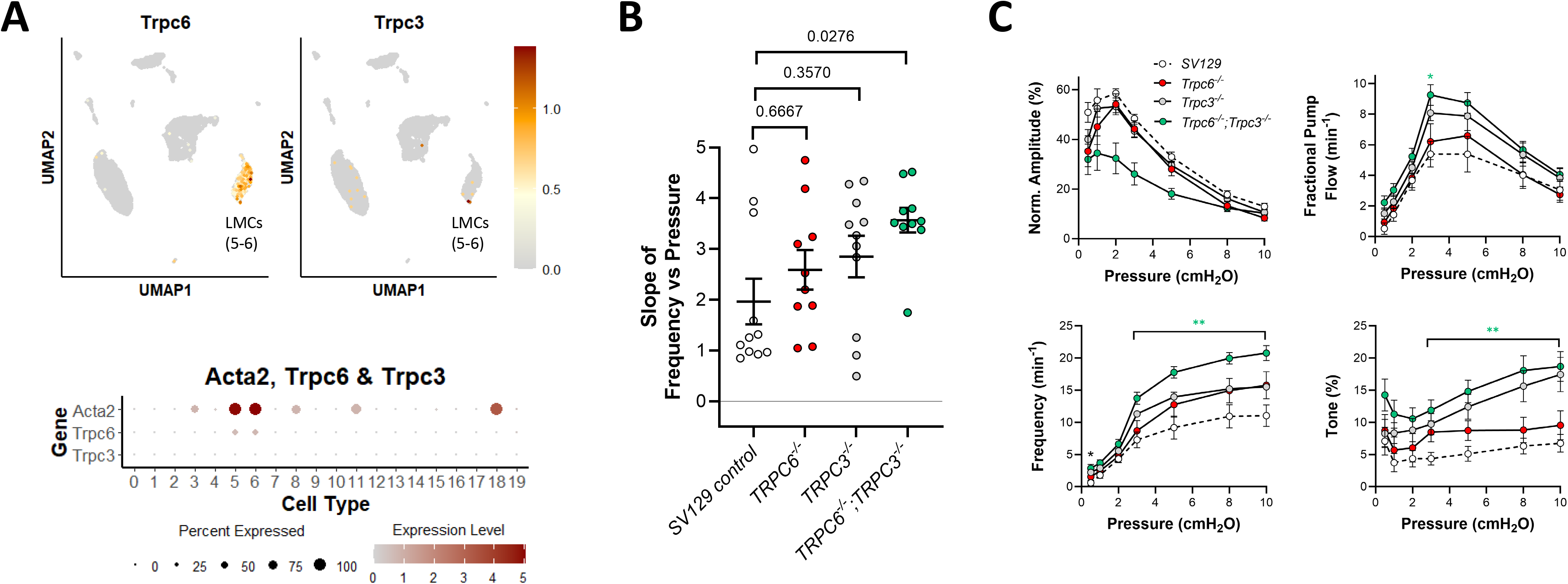
Consequences of *Trpc6* and/or *Trpc3* deletion. **A**) UMAP and bubble plots of *Trpc6 and Trpc3* expression in the various IALV cell clusters. **B**) Slope of F-P relationship for popliteal lymphatics from SV129 controls, *Trpc6^-/-^, Trpc3^-/-^ and Trpc6^-/-^;Trpc3^-/-^* mice. **C**) Summary plots of contraction parameters as a function of pressure for the 4 respective groups of vessels. SV129 control N=8, n=11; *Trpc6^-/-^* N=5, n=10; *Trpc3^-/-^* N=7, n=11; *Trpc6^-/-^;Trpc3^-/-^* N=5, n=10.

### Role of TRPM4

Multiple studies have implicated TRPM4 channels in the arterial myogenic response (Gonzales & Earley, 2013; Earley & Brayden, 2015; Zhu *et al*., 2026), but to date no published studies have examined the role of TRPM4 in lymphatic vessel contractility. scRNAseq analysis revealed that *Trpm4* was expressed at modest levels in only ∼10% of LMCs, and in a subset of LECs, but at higher levels in some AdvCs (**Fig. 6A**). We tested the role of TRPM4 in pressure-induced lymphatic chronotropy by comparing the responses of Myh11-CreER^T2^*;Trpm4^f/f^* (*Trpm4* smKO) vessels to *Trpm4^f/f^* vessels. The lack of significant reductions in F-P slope of *Trpm4* smKO vessels (**Fig. 6B**) suggests that TRPM4 is not required for pressure-induced lymphatic chronotropy. Contraction data comparing *Trpm4* smKO vessels to *Trpm4^f/f^* popliteal vessels are shown in **Fig. 6C**. There were no significant differences between the contraction parameters of *Trpm4* smKO and *Trpm4^f/f^* vessels at any pressure.

**Figure 6.**
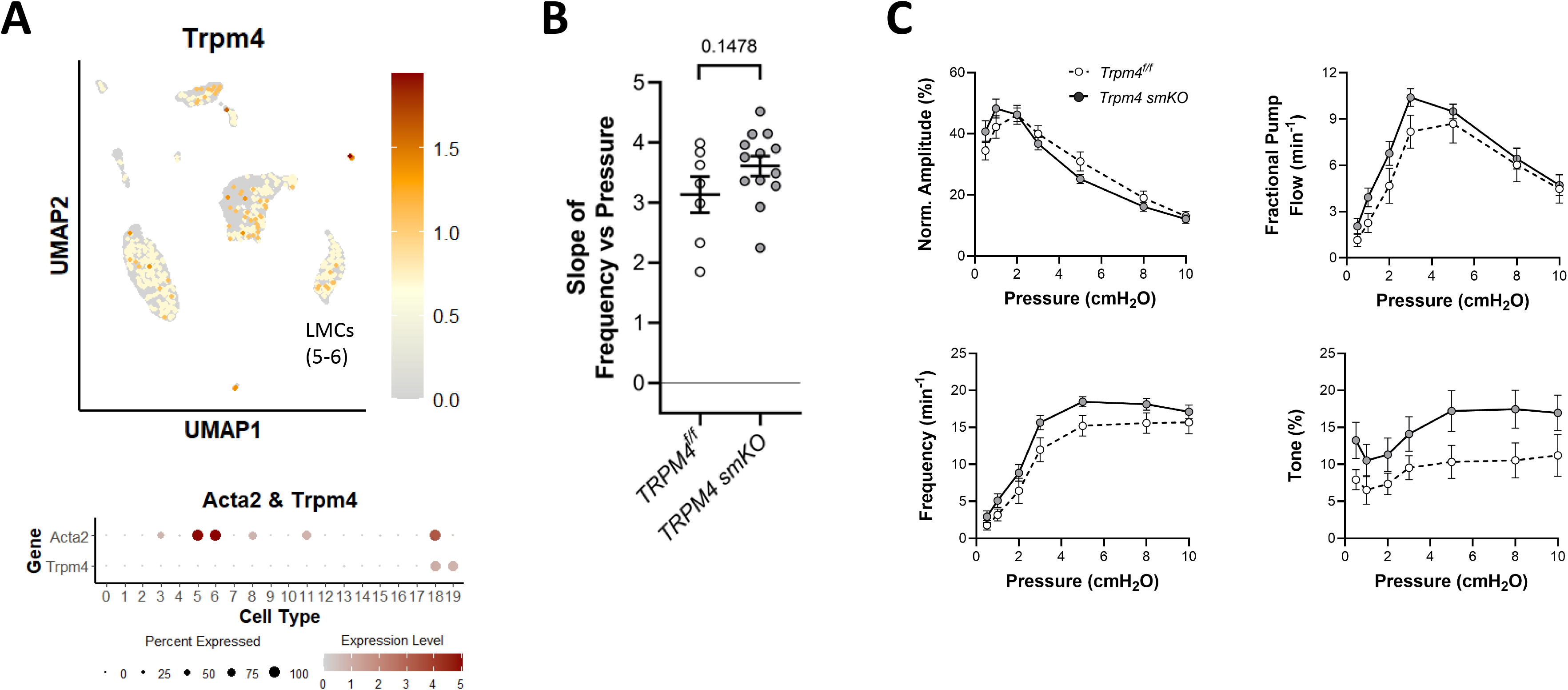
Consequences of *Trpm4* deletion. **A**) UMAP and bubble plots of *Trpm4* expression in the various IALV cell clusters. **B**) Slope of F-P relationship for popliteal lymphatics from *Trpm4^f/f^ and Trpm4* smKO mice. **C**) Summary plots of contraction parameters as a function of pressure for the two respective groups of vessels. *Trpm4* smKO N=6, n=13; *Trpm4^f/f^* N=4, n=7.

### Role of PKD2

Studies by Jaggar and colleagues indicate that SM-specific deletion of *Pkd2* attenuates pressure-induced depolarization and myogenic tone in rat pial arteries (Narayanan *et al*., 2013) and blunts pressure-induced constriction in hindlimb arteries (Bulley *et al*., 2018). The roles of PKD1/2 channels in lymphatic vessel contractility are not known. scRNAseq analysis revealed *Pkd2* and *Pkd1* expression in most LMCs and AdvCs (**Fig. 7A**). We tested the possible involvement of PKD2 channels in pressure-induced lymphatic chronotropy by comparing the responses of Myh11-CreER^T2^*;Pkd2^f/f^*(*Pkd2* smKO) vessels to *Pkd2^f/f^* vessels (**Fig. 7B**). The lack of a significant reduction in the F-P slope for *Pkd2* smKO vessels suggests that PKD2 is not required for pressure-induced lymphatic chronotropy. Contraction data comparing *Pkd2* smKO vessels to *Pkd2^f/f^*popliteal vessels are shown in **Fig. 7C**. There were no significant differences between the contraction parameters of *Pkd2* smKO and *Pkd2^f/f^* vessels at any pressure.

**Figure 7.**
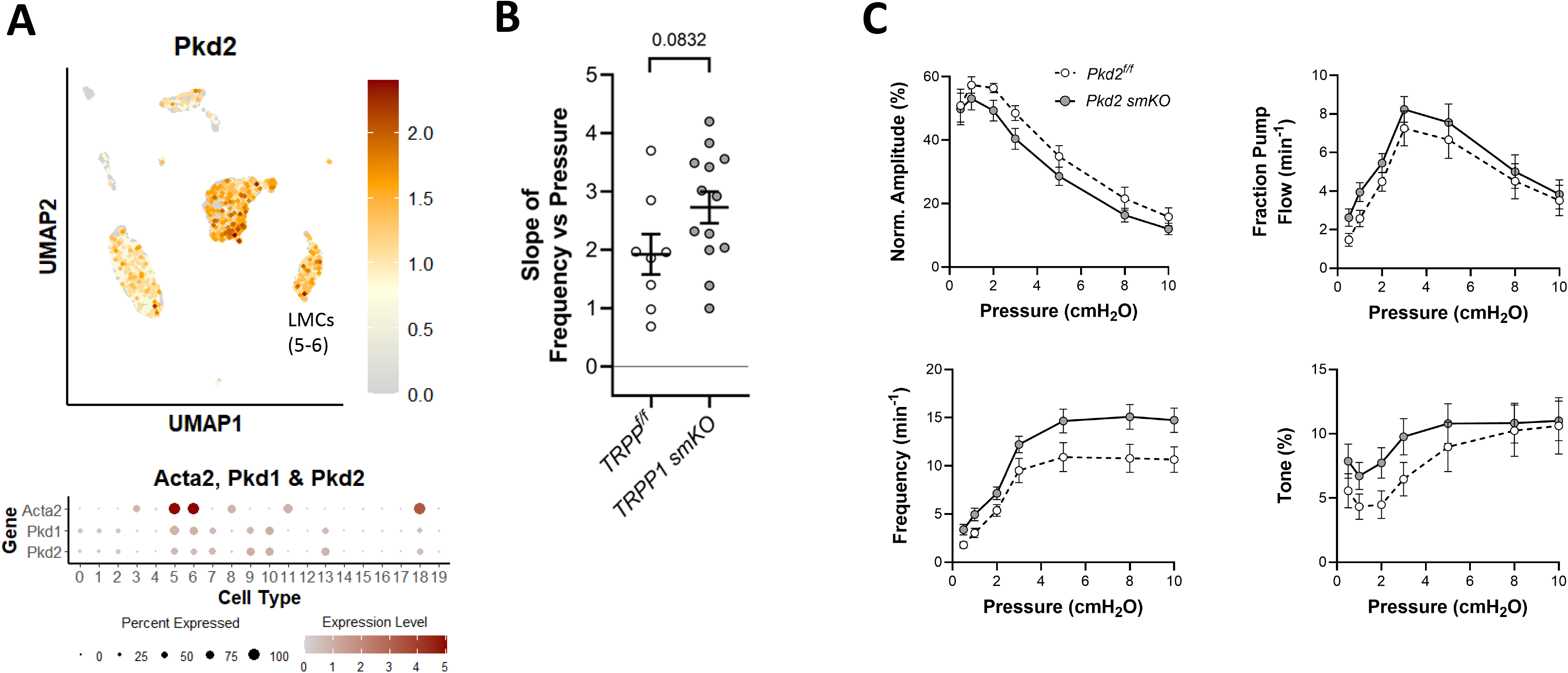
Consequences of *Pkd2* (TRPP1) deletion. UMAP and bubble plots of *Pkd2* expression in the various IALV cell clusters. **B**) Slope of F-P relationship for popliteal lymphatics from *Pkd2 ^f/f^ and Pkd2* smKO mice. **C**) Summary plots of contraction parameters as a function of pressure for the two respective groups of vessels. *Pkd2* smKO N=6, n=13; *Pkd2 ^f/f^* N=4, n=8.

### Role of TRPV4

In the blood vasculature, TPRV4 has been studied extensively in the context of Ca^2+^ influx pathways in endothelium (Chen & Sonkusare, 2020). However, TRPV4 is also expressed in VSMCs of some arteries (Mercado *et al*., 2014) and one report indicates that pharmacological inhibition of TRPV4 channels blunts pressure-induced VSMC depolarization and myogenic vasoconstriction of preglomerular arteries from neonatal pigs (Soni *et al*., 2017). Our scRNAseq analysis revealed that *Trpv4* was largely undetectable in LMCs but expressed at higher levels in almost all LECs, macrophages and some AdvCs (**Fig. 8A**). Nonetheless, we tested the role of TRPV4 channels in pressure-induced lymphatic chronotropy by comparing the responses of Myh11-CreER^T2^*;Trpv4^f/f^* (*Trpv4* smKO) vessels to *Trpv4^f/f^* vessels. The lack of a significant reduction in the F-P slope for *Trpv4* smKO vessels suggests that TRPV4 is not required for pressure-induced lymphatic chronotropy (**Fig. 8B**). Contraction data comparing *Trpv4* smKO and *Trpv4^f/f^* popliteal vessels are shown in **Fig. 8C**. Contraction amplitude was impaired at several pressures in *Trpv4* smKO vessels, suggesting some involvement of TRPV4 channels in control of contractile strength, possibly as a result of macrophage-derived products (Schulz *et al*., 2025). Frequency was significantly different at two lower pressures in *Trpv4* smKO vessels (**Fig. 8C**), where it was *elevated* rather than reduced. We also tested the F-P relationship in *Trpv4^-/-^* (global KO) vessels and confirmed that there was no significant impairment in pressure-induced lymphatic chronotropy (average slope = 2.7 ± 0.2; n=18) compared to WT vessels (average slope = 2.8 ± 0.3; data not shown). Collectively, these results indicate that TRPV4 channels are not required for pressure-induced lymphatic chronotropy.

**Figure 8.**
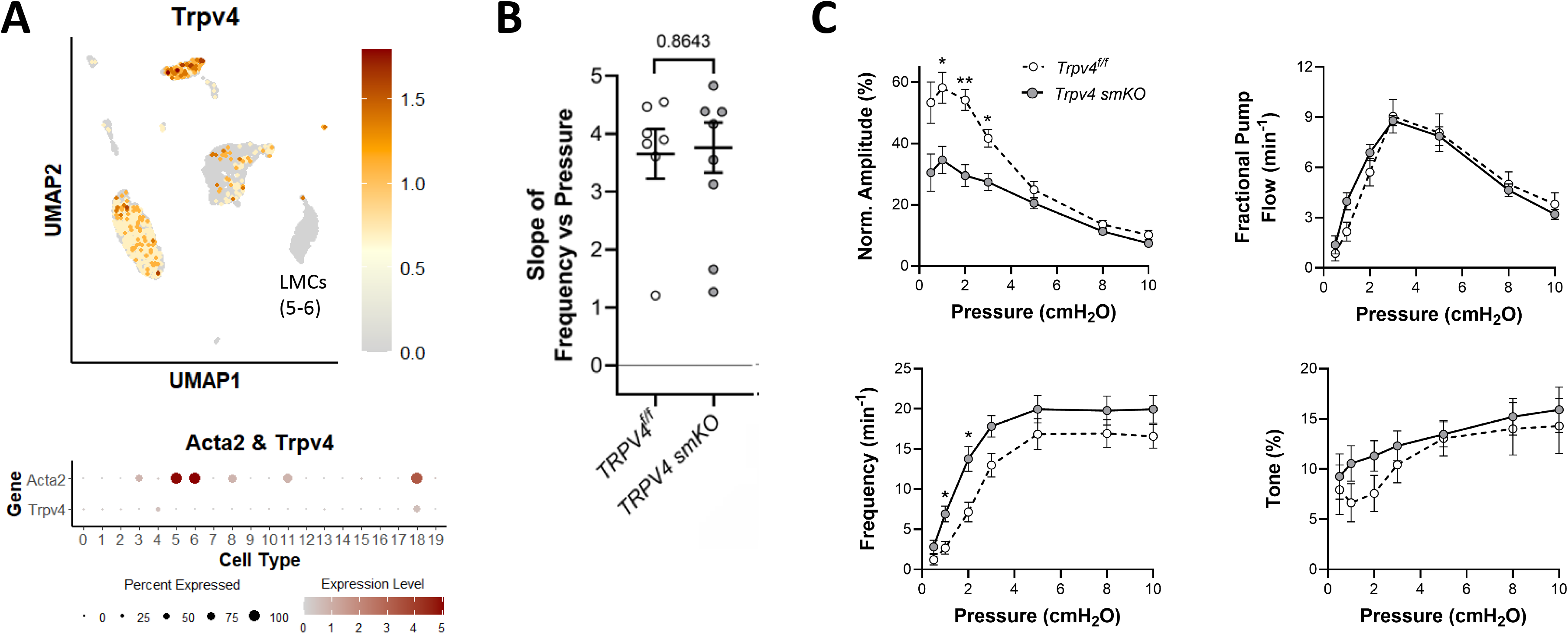
Consequences of *Trpv4* deletion. UMAP and bubble plots of *Trpv4* expression in the various IALV cell clusters. **B**) Slope of F-P relationship for popliteal lymphatics from *Trpv4^f/f^ and Trpv4* smKO mice. **C**) Summary plots of contraction parameters as a function of pressure for the two respective groups of vessels. *Trpv4* smKO N=5, n=10; *Trpv4^f/f^* N=3, n=7.

### Role of TRPV2

TRPV2 is an osmosensitive cation channel (Nilius & Droogmans, 2001). Down-regulation of *Trpv2* expression in mouse aortic VSMCs reduced the amplitude of cation currents activated by hypotonic swelling (Muraki *et al*., 2003). In another study, application of a hypotonic bath solution stimulated TRPV2-like whole-cell cation currents and Ca^2+^ influx in VSMCs isolated from rat retinal arteries—responses that were attenuated by the TRPV2 inhibitor tranilast (McGahon *et al*., 2016). Additionally, the administration of tranilast to rat retinal arteries resulted in loss of pre-existing myogenic tone whereas pre-incubation with a TRPV2 blocking antibody prevented the development of myogenic tone (McGahon *et al*., 2016). Our scRNAseq analysis revealed that *Trpv2* was highly expressed in 63% of *Acta2*-expressing LMCs (**Fig. 9A**). As we did not have access to *Trpv2* null or floxed mice, we tested the potential role of TRPV2 channels in pressure-induced lymphatic chronotropy by bath application of tranilast (100 μM) to WT popliteal lymphatic vessels. There was no significant reduction in the F-P slope for tranilast-treated vessels (**Fig. 9B**). We also tested the effects of SET2, a recently reported TRPV2 inhibitor with a lower IC_50_ (460 nM) than tranilast. The application of SET2 (1 μM) caused a transient cessation of spontaneous contractions for 2-5 min in 5 of 6 vessels, but the contractions returned within 10 min in the continued presence of SET2. Subsequent pressure steps revealed that SET2 did not significantly alter the F-P slope (**Fig. 9B**). Contraction data comparing WT and tranilast- and SET2-treated WT popliteal vessels are shown in **Fig. 9C**. These results suggest that TRPV2 is not required for pressure-induced lymphatic chronotropy.

**Figure 9.**
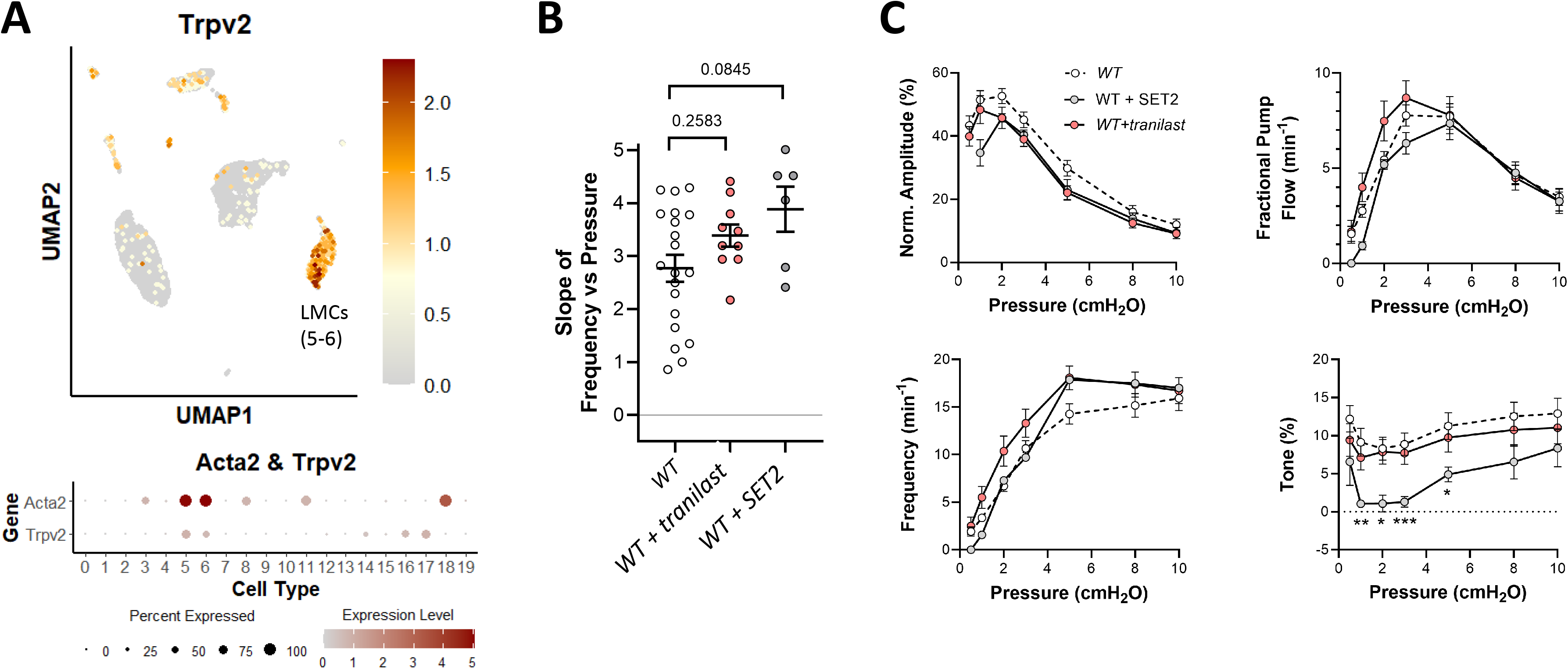
Consequences of Trpv2 inhibition. UMAP and bubble plots of *Trpv2* expression in the various IALV cell clusters. **B**) Slope of F-P relationship for popliteal lymphatics from WT mice with/without treatment with the TRPV2 inhibitors tranilast (10 μM) and SET2 (1 μM). **C**) Summary plots of contraction parameters as a function of pressure for the three respective groups of vessels. WT+tranilast N=5, n=10; WT+SET2 N=3, n=6.

### Role of ENaC

The epithelial Na^+^ channel (ENaC), comprised of a heterotrimer under the expression of the genes *Scnn1a*, *Scnn1b*, *Scnn1g*, *Scnn1d*, is expressed in both the endothelial and smooth muscle layers of arteries. αENaC is the principal isoform in the endothelium and studies of native epithelial or heterologous channels reconstituted into bilayers suggest that αENaC is intrinsically mechanosensitive (Price *et al*., 2000; Benos, 2004). Work from the Drummond laboratory suggests that the βENaC isoform is expressed in VSMCs and plays a critically important role in myogenic constriction (Drummond, 2012). However, our scRNAseq analysis revealed almost no expression of αENaC subunits in LMCs and only weak expression in the LEC population (**Fig. 10A**); message for βENaC was essentially undetectable. Nevertheless, we tested the possible role of ENaC in pressure-induced lymphatic chronotropy by administering two widely used ENaC inhibitors, amiloride and benzamil, to WT popliteal lymphatic vessels. The data in **Fig. 10B** show that neither amiloride nor benzamil, at concentrations known to inhibit arterial myogenic constriction [5 and 1 μM, respectively (Jernigan & Drummond, 2006)], caused significant impairment in pressure-induced lymphatic chronotropy. Contraction data for amiloride- and benzamil-treated WT popliteal vessels are shown in **Fig. 10C**. Both ENaC inhibitors tended to attenuate contraction amplitude to some degree (significant only for amiloride), with benzamil also significantly increasing tone at almost all pressure levels. However, the F-P relationships were nearly identical between WT and amiloride- or benzamil-treated vessels, reinforcing the conclusion that ENaC is not involved in pressure-induced lymphatic chronotropy.

**Figure 10.**
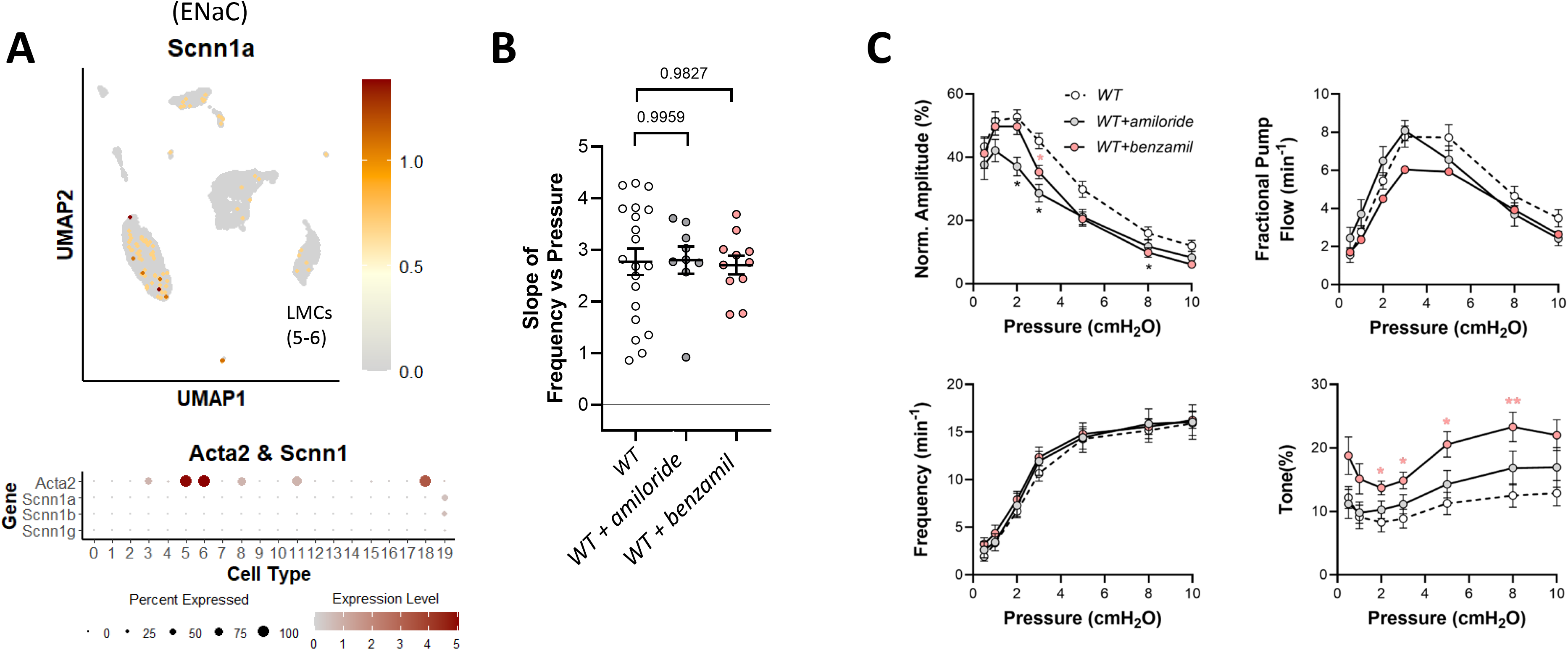
Consequences of ENaC inhibition. UMAP and bubble plots of *ENaC* subunit expression in the various IALV cell clusters. **B**) Slope of F-P relationship for popliteal lymphatics from WT mice with/without treatment with the ENaC inhibitors amiloride (5 μM) and benzamil (1 μM). WT N=15, n=20, WT+amiloride N=5, n=9; WT+benzamil N=6, n=11.

### Role of NCX1

The sodium-calcium exchanger (NCX) has a well-documented role in the arterial myogenic response (Zhang, 2013) and in controlling the pacemaker activity of sinoatrial nodal cells, where it couples SR calcium oscillations caused by calcium movement through ryanodine channels (Vinogradova *et al*., 2004; Vinogradova *et al*., 2005), termed local calcium releases (LCRs), to AP ignition. LCRs may occur throughout diastole (Monfredi *et al*., 2013), but their role is particularly critical in late diastole where they activate NCX and the ensuing addition of inward current, plus positive feedback of further calcium influx through L-type channels, provides sufficient depolarization to cross the voltage threshold for initiating AP firing (Bogdanov *et al*., 2001; Lyashkov *et al*., 2018b). This process is possibly analogous to the electrophysiological pacemaking mechanism in LMCs and, indeed, a previous study demonstrated a functional NCX1 exchanger in human lymphatic muscle (Telinius *et al*., 2015). scRNAseq analysis revealed moderate expression of *Slc8a1*, which encodes NCX1, in >80% of LMCs and stronger expression in LECs and some immune cells (**Fig. 11A**). We tested the role of NCX1 in pressure-induced chronotropy by comparing the responses of Myh11-CreER^T2^*;Slc8a1^f/f^* (*Slc8a1* smKO) and *Slc8a1^f/f^*vessels. The lack of a significant reduction in the F-P slope for *Slc8a1* smKO vessels suggests that NCX1 is not required for pressure-induced lymphatic chronotropy (**Fig. 11B**). Contraction data for *Slc8a1* smKO and *Slc8a1^f/f^* popliteal vessels are shown in **Fig. 11C** and reveal no significant differences between the contraction parameters of *Slc8a1* smKO and *Slc8a1^f/f^* vessels.

**Figure 11.**
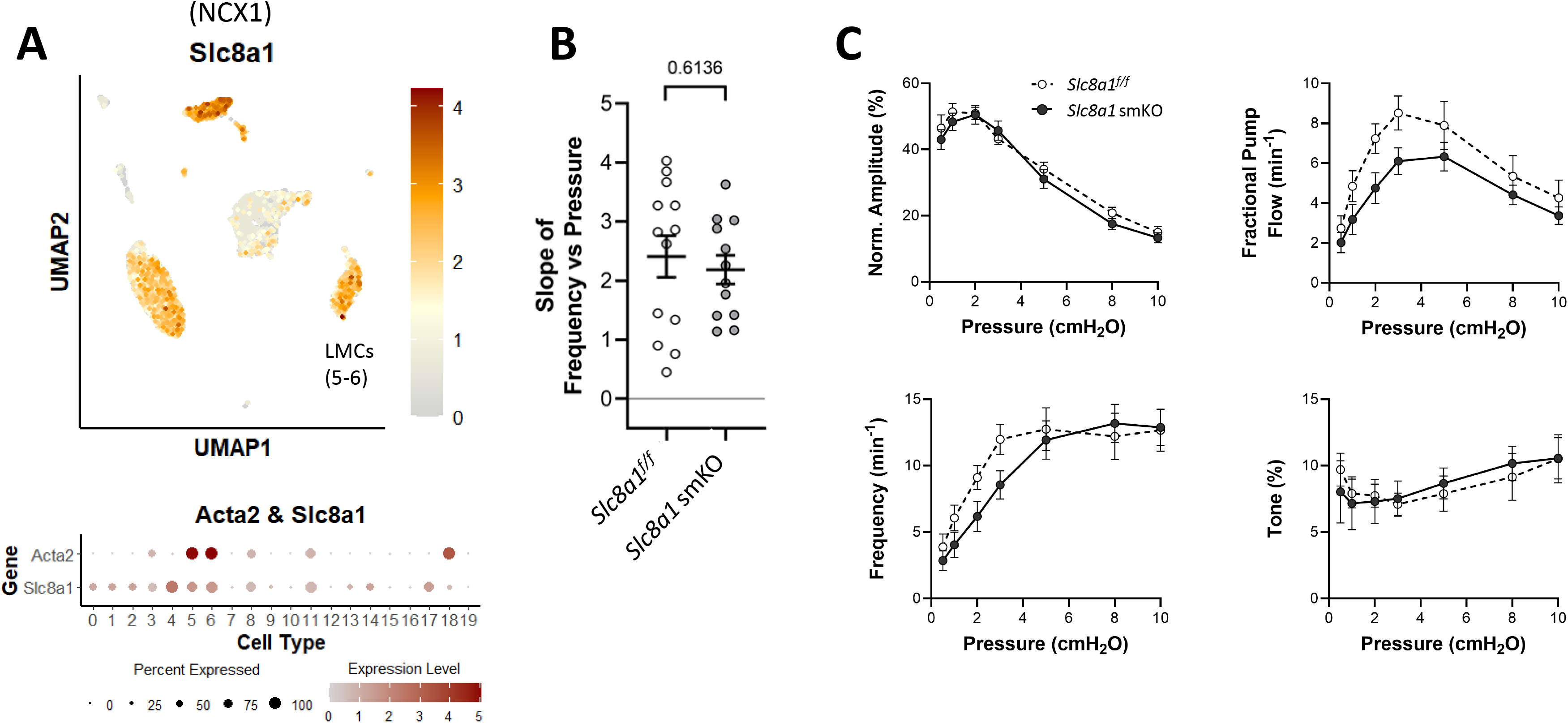
Consequences of *Slc8a1* (NCX1) deletion. UMAP and bubble plots of *Slc8a1* expression in the various IALV cell clusters. **B**) Slope of F-P relationship for popliteal lymphatics from *Slc8a1^f/f^ and Slc8a1* smKO mice. **C**) Summary plots of contraction parameters as a function of pressure for the two respective groups of vessels. *Slc8a1^f/f^* N=7, n=12; *Slc8a1* smKO N=7, n=13.

### Role of HCN

HCN (Hyperpolarization-activated cyclic-nucleotide-gated) channels contribute an inward “funny” current (I_f_) to the diastolic pacemaking potential of sinoatrial node cells (Accili *et al*., 2002) and are also involved in the intrinsic rhythmicity of bladder and uterine smooth muscle (Alotaibi *et al*., 2017; Mader *et al*., 2018). In LMCs they could potentially contribute to diastolic depolarization. The HCN blockers CsCl, ZD7288, and ivabradine all lower the rate of spontaneous contractions of lymphatic vessels in rat diaphragm (Negrini *et al*., 2016), but inhibition requires substantially higher concentrations than the documented IC_50_ values for HCN channel blockade [see discussion in (Davis & Zawieja, 2024)]. In contrast, all three inhibitors *increase* the spontaneous contraction frequency of human lymphatic vessels at concentrations near or slightly higher than the IC_50_ values for HCN inhibition (Majgaard *et al*., 2022), suggesting off-target effects in some lymphatic vessels. Our scRNAseq analysis revealed essentially no expression of any of the four HCN isoforms in LMCs (**Fig. 12A**), however, it is possible that the expression was too low to detect without deep sequencing. Due to the proven role of HCN channels in cardiac pacemaking, we tested the role of HCN channels in pressure-induced lymphatic chronotropy of WT mouse popliteal lymphatics using the HCN inhibitor ivabradine (at a concentration of 3 μM, which is slightly higher than the IC_50_ of 2.5 μM), and the more effective inhibitor, zatebradine, at a concentration of 3 μM (IC_50_ = 480 nM). The lack of significant reductions in the F-P slope for ivabradine- or zatebradine-treated WT vessels suggests that HCN channels are not required for pressure-induced lymphatic chronotropy (**Fig. 12B**). Contraction data for ivabradine- and zatebradine-treated WT vessels are shown in **Fig. 12C** and reveal no substantial differences from WT vessels, except for a single pressure at which each HCN channel inhibitor actually *increased* the contraction frequency. Collectively, these results recapitulate the findings of the in-depth study by Majgaard et al (Majgaard *et al*., 2022) and reinforce the conclusion that HCN channels are not required for pressure-induced lymphatic chronotropy.

**Figure 12.**
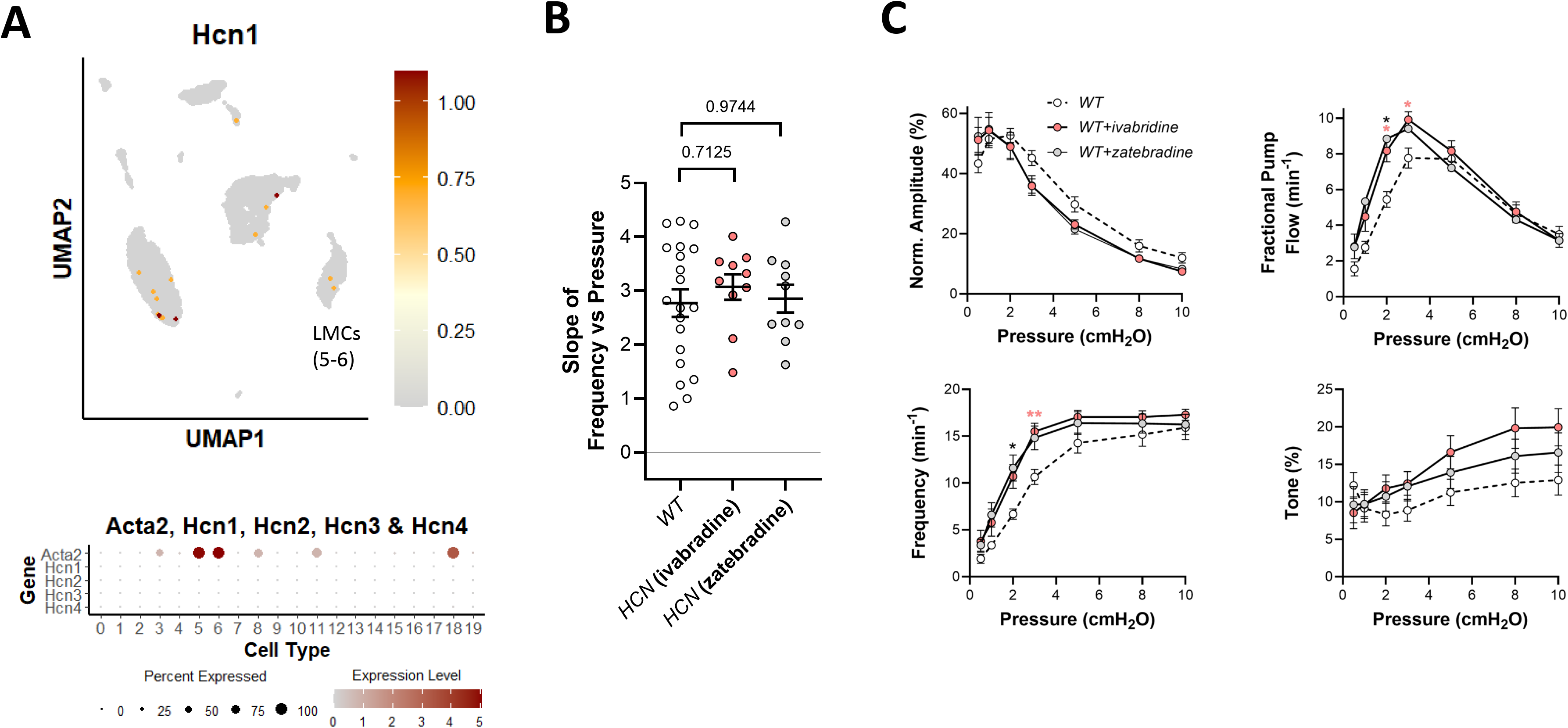
Consequences of HCN inhibition. UMAP and bubble plots of *Hcn gene* family expression in the various IALV cell clusters. **B**) Slope of F-P relationship for popliteal lymphatics from WT mice with/without treatment with the HCN inhibitors ivabradine (3 μM) and zatebradine (3 μM). WT N=15, n=20, WT+zatebradine. N=5, n=10; WT+ivabradine N=5, n=10.

### Role of Kv7.4

Kv7 channels are activated at more negative membrane potentials than Kv1-family delayed rectifier K^+^ channels and thus are implicated in control of VSMC excitability near the resting membrane potential (Yeung & Greenwood, 2005; Zhong *et al*., 2010; Mani & Byron, 2011). The Kv7 blocker, XE991, enhances myogenic tone in isolated mesenteric and renal arteries and hypotonic swelling suppresses an XE991-sensitive Kv current in patch-clamped VSMCs (Schleifenbaum *et al*., 2014). Collectively, these results imply that mechanosensitive *inhibition* of Kv7.4 may increase VSMC excitability. The effects of Kv7 channel inhibition on lymphatic function are unknown. Our scRNAseq analysis revealed moderately strong expression of *Kcnq4* in >50% LMCs and in some AdvCs (**Fig. 13A**). We tested whether pressure-induced lymphatic chronotropy was altered in Myh11-CreER^T2^*;Kcnq4^f/f^* (*Kcnq4* smKO) mice. However, the lack of a significant reduction in the F-P slope for *Kcnq4* smKO vessels suggests that Kv7.4 channels are not required for pressure-induced lymphatic chronotropy (**Fig. 13B**). Contraction data for *Kcnq4* smKO and WT vessels are compared in **Fig. 13C**. There were no significant differences in any of the parameters at any pressure.

**Figure 13.**
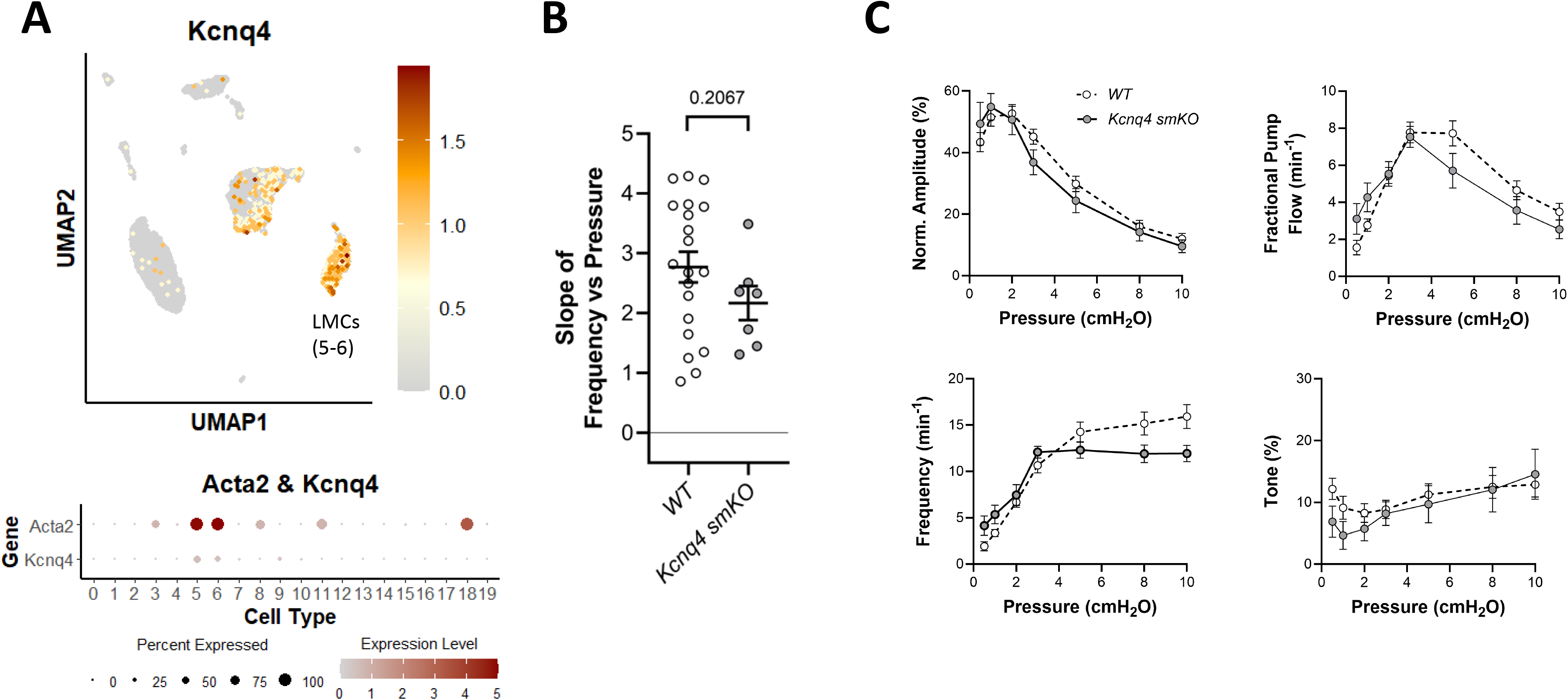
Consequences of *Kcnq4* deletion. UMAP and bubble plots of *Kcnq4* expression in the various IALV cell clusters. **B**) Slope of F-P relationship for popliteal lymphatics from WT *and Kcnq4* smKO mice. **C**) Summary plots of contraction parameters as a function of pressure for the two respective groups of vessels. *Kcnq4* smKO N=5, n=7.

### Role of Kir channels

Inwardly rectifying K^+^ channels, including Kir2.1 and Kir2.2, encoded by *Kcnj2* and *Kcnj12* respectively, have been implicated in shear-stress-induced responses of arterial endothelium (Ahn *et al*., 2017) and swelling-induced currents in arterial smooth muscle cells (Sancho *et al*., 2019). However, their potential roles in controlling LMC membrane potential and pressure-induced lymphatic chronotropy are unknown. Our scRNAseq analysis revealed modest expression of *Kcnj8* (Kir6.1), the inward rectifier which forms the pore forming unit of the K_ATP_ channel, in >50% LMCs (**Fig. 14A**), but essentially no expression of *Kcnj2* or *Kcnj12* in LMCs. We previously showed that deletion of *Kcnj8* does not alter the F-P relationship in popliteal lymphatics (Davis *et al*., 2020). Here, we tested the effects of BaCl_2_, a selective inhibitor of Kir2.1/2.2 at a concentration of 100 μM (Longden *et al*., 2017; Sancho *et al*., 2017; Sancho *et al*., 2019), on the F-P relationship of WT popliteal lymphatic vessels. Kir inhibition resulted in a significantly reduced slope of the F-P relationship compared to WT vessels (**Fig. 14B**), but the consistent effect of BaCl_2_ was to flatten the slope of the F-P relationship by substantially *increasing* the contraction frequency at low pressures while only slightly *increasing* the contraction frequency at elevated pressures (**Fig. 14C**). This contrasts with the effect of ANO1 inhibition or deletion: in the absence of ANO1 activity, the contraction frequency was significantly *reduced* at all pressures, particularly at low pressures (**Fig. 2C**). Thus, the inhibition of Kir channels does impair pressure-induced chronotropy but does so by differentially *elevating* contraction frequency as a function of pressure.

**Figure 14.**
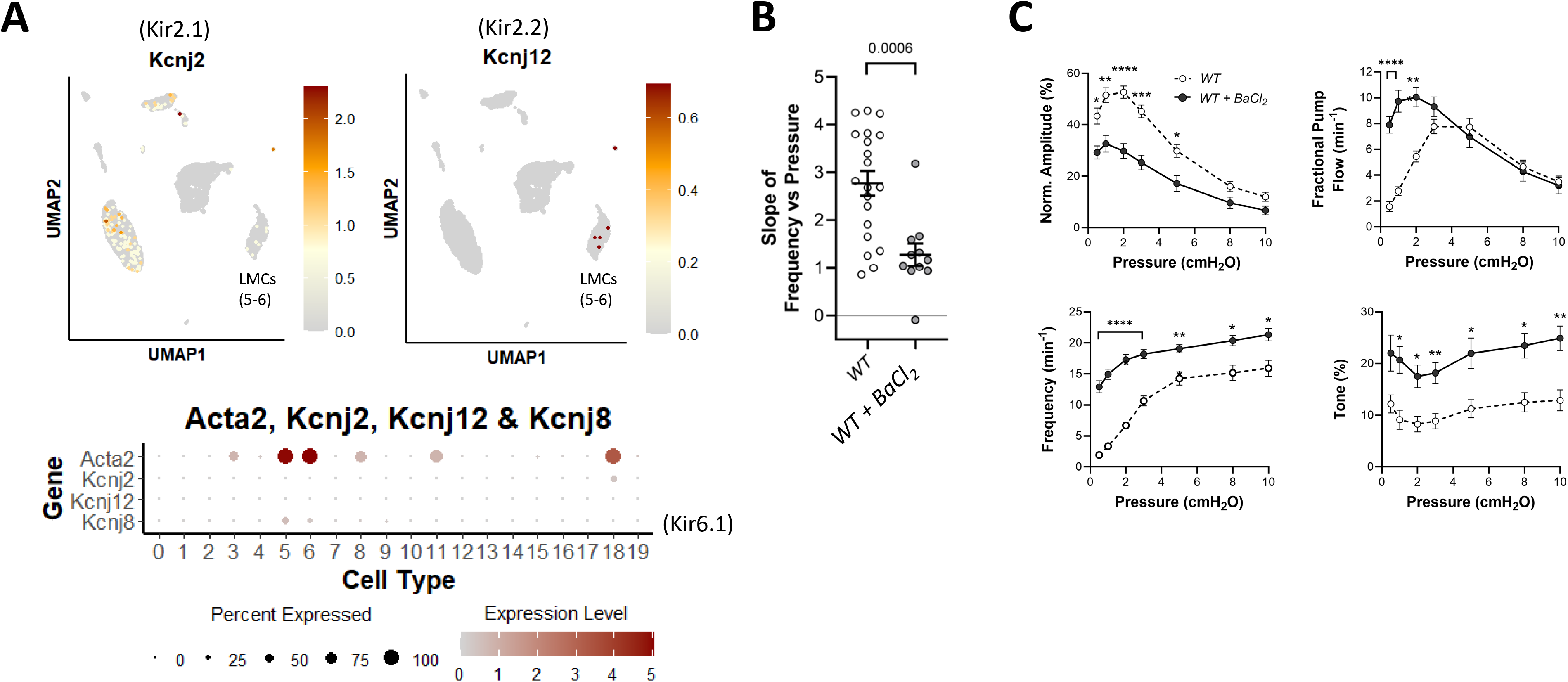
Consequences of Kir2.x inhibition. UMAP and bubble plots of *Kcnj2* (Kir2.1*), Kcnj12 (*Kir2.2) and *Kcnj8* (*Kir6.1)* expression in various IALV cell clusters. **B**) Slope of F-P relationship for popliteal lymphatics from WT mice with/without treatment with the Kir2 inhibitor BaCl_2_ (100 μM). WT N=15, n=20, WT+BaCl_2_ N=4, n=11.

### Roles of L- and T-type Ca^2+^ channels

The L-type, voltage-gated Ca^2+^ channel (Cav1.2) carries the majority of current contributing to the action potential in LMCs (van Helden, 1993; To *et al*., 2020). scRNAseq analysis confirmed that >90% of LMCs expressed message for the pore forming subunit of Cav1.2 (*Cacna1c*), along with other accessory subunits of the channel responsible for gating properties (α2δ) or membrane targeting (β) (**Fig. 15A**). SM-specific knockout of *Cacna1*, using either the Myh11-CreER^T2^ or Itga8-CreER^T2^, eliminated essentially all propulsive contractions from popliteal lymphatic vessels (Warthi *et al*., 2022; Davis *et al*., 2023e). Residual activity in some vessels was limited to small, localized diameter oscillations (<5 μm in amplitude) with irregular frequencies that were largely independent of pressure. As expected, the slope of the F-P relationship for Myh11-CreER^T2^;*Cacna1c^f/f^*(*Cacna1c* smKO) vessels was significantly reduced compared to *Cacna1c^f/f^*control vessels (**Fig. 15B**) and contraction frequencies were significantly suppressed compared to control vessels at all pressures (**Fig. 15C**).

**Figure 15.**
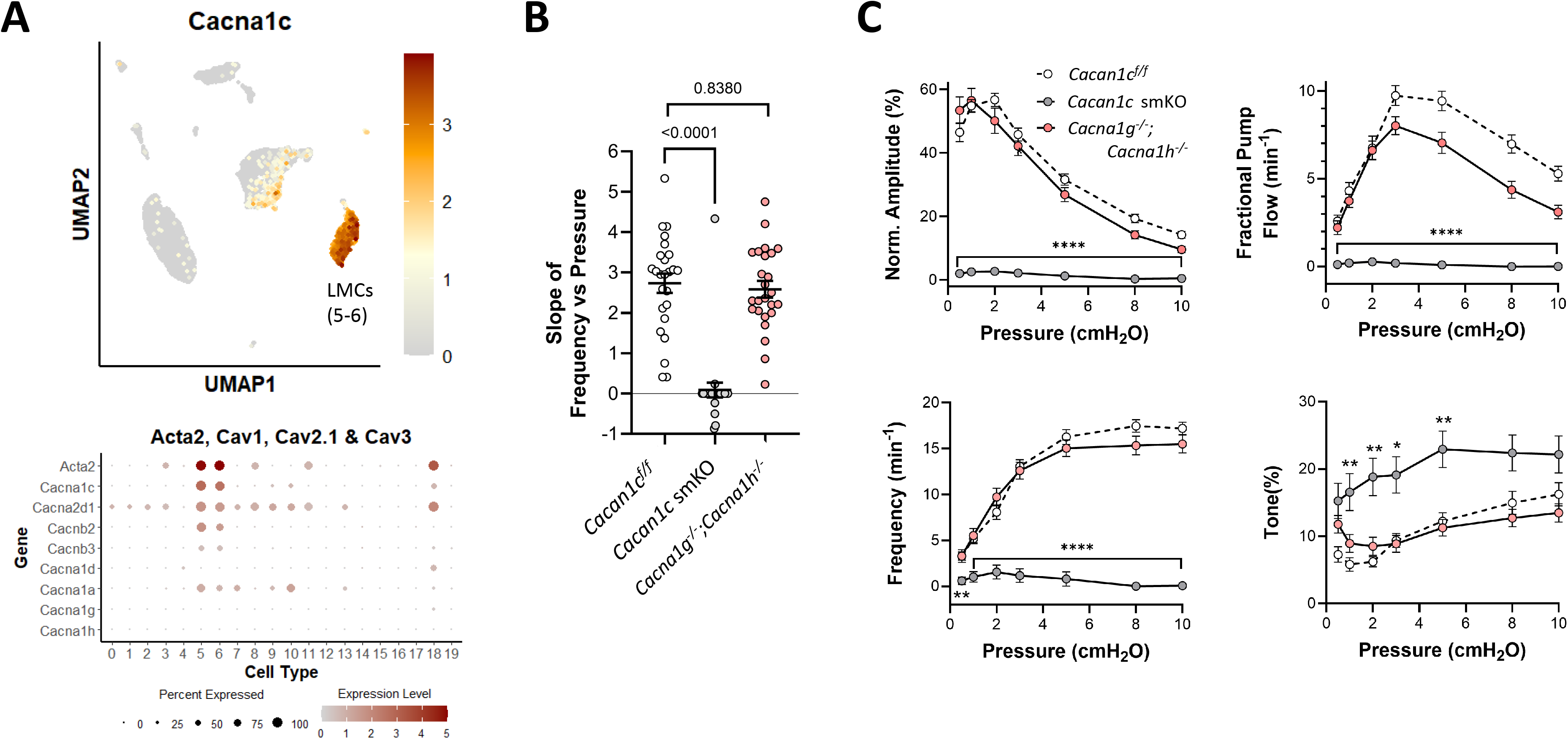
Consequences of *Cacna1c* (Cav1.2) and *Cacna1g/Cacna1h* (Cav3.1/3.2) deletion. UMAP and bubble plots of the expression of voltage-gated Ca^2+^ channels and their accessory subunits in the various IALV cell clusters. *Cacna1c*: alpha-1c, *Cacna2d1*: α2/δ1, *Cacnb2*: β2, *Cacnb3*: β3, *Cacna1d*: alpha-1d (Ca_v_1.3), *Cacna1a*: Ca_v_2.1 (P/Q), *Cacna1g*: Ca_v_3.1, *Cacna1h*: Ca_v_3.2. **B**) Slope of F-P relationship for popliteal lymphatics from *Cacan1c^f/f^, Cacan1c* smKO and *Cacna1g^-/-^;Cacna1h^-/-^* mice. **C**) Summary plots of contraction parameters as a function of pressure for the three respective groups of vessels. Horizontal bar indicates range of pressures with same specified statistical significance. *Cacna1c^f/f^* N=10, n=20; *Cacna1c* smKO N=14, n=24; *Cacna1g^-/-^;Cacna1h^-/-^* N=16, n=25.

The T-type Ca^2+^ channel (Cav3.x) is critical for pacemaking in SA node (Lyashkov *et al*., 2018a), GI smooth muscle (Sanders, 2019), and renal pelvis smooth muscle (Grainger *et al*., 2022). Two Cav3 isoforms are expressed in LMCs: *Cacna1g* and *Cacna1h,* encoding Cav3.1 and Cav3.2 respectively (Lee *et al*., 2014; To *et al*., 2020). One report suggests that Cav3 channels are selectively involved in regulating the contraction frequency (but not amplitude) of rat mesenteric lymphatic collectors (Lee *et al*., 2014). We previously tested the roles of Cav3.1 and Cav3.2 in mouse lymphatic collectors and isolated LMCs after generating *Cacna1g^-/-^;Cacna1h^-/-^* double KO mice (To *et al*., 2020). Here, we analyzed the slope of the F-P relationship in popliteal lymphatics from *Cacna1g^-/-^;Cacna1h^-/-^*mice but found no significant differences compared to either WT or *Cacan1c^f/f^* control vessels (**Fig. 15B**), indicating that Cav3 channels are not involved in determining the F-P relationship. No significant differences in frequency were noted between *Cacna1g^-/-^ ;Cacna1h^-/-^* and control vessels at any pressure (**Fig. 15C**).

### Upstream pathways mediating pressure-induced chronotropy

Having established that ANO1, out of all the cation and K^+^ channels tested, is the only channel mediating pressure-induced lymphatic chronotropy, we then turned to an investigation of mechanosensitive pathways upstream from ANO1 channel activation.

### Role of IP_3_R in pressure-induced chronotropy

Calcium release as a consequence of IP_3_ generation is known to activate a number of ion channels and other proteins, including ANO1. We recently showed that genetic deletion of *Itpr1* (encoding IP_3_R1) from mouse IALVs results in a reduction in spontaneous contraction frequency and a blunting of pressure-induced chronotropy similar to what is observed after *Ano1* deletion (Zawieja *et al*., 2023). scRNAseq analysis confirmed expression of *Itpr1* in >50% of LMCs and only very weak expression of *Itpr2* and *Itpr3* (**Fig. 16A**). Here, we examined the consequences of *Itpr1* deletion in popliteal lymphatics by comparing the F-P relationship for vessels from Myh11-CreER^T2^*;Itpr1^f/f^*(*Itpr1* smKO) and *Itpr1^f/f^* mice. SM-specific deletion of *Itpr1* resulted in a significant reduction in the slope of the F-P relationship from 2.6 ± 0.3 to 0.6 ± 0.1 (**Fig. 16B**). Contraction data for *Itpr1* smKO and *Itpr1^f/f^*control vessels are shown in **Fig. 16C**. The most striking features are 1) the significant reductions in frequency at all pressures and 2) a significant reduction in tone at elevated pressures in *Itpr1* smKO vessels. The latter effect contrasts with the *increases* in tone observed after ANO1 inhibition or deletion (**Fig. 2C**). Collectively, these results show that IP_3_R1 is critically important for pressure-induced lymphatic chronotropy in popliteal collecting vessels in much the same way that was previously documented in IALVs. Deletion of *Itpr1* in LMCs produces an effect on pressure-induced chronotropy similar to that of ANO1 deletion. Examples of the contraction patterns observed in *Itpr1^f/f^* and *Itpr1* smKO vessels are shown in **Figure 17**; in the KO vessel the basal contraction frequency was only 1.8 min^-1^ at P=3 cmH_2_O and increased to only 1.9 min^-1^ at P=5 cmH_2_O.

**Figure 16.**
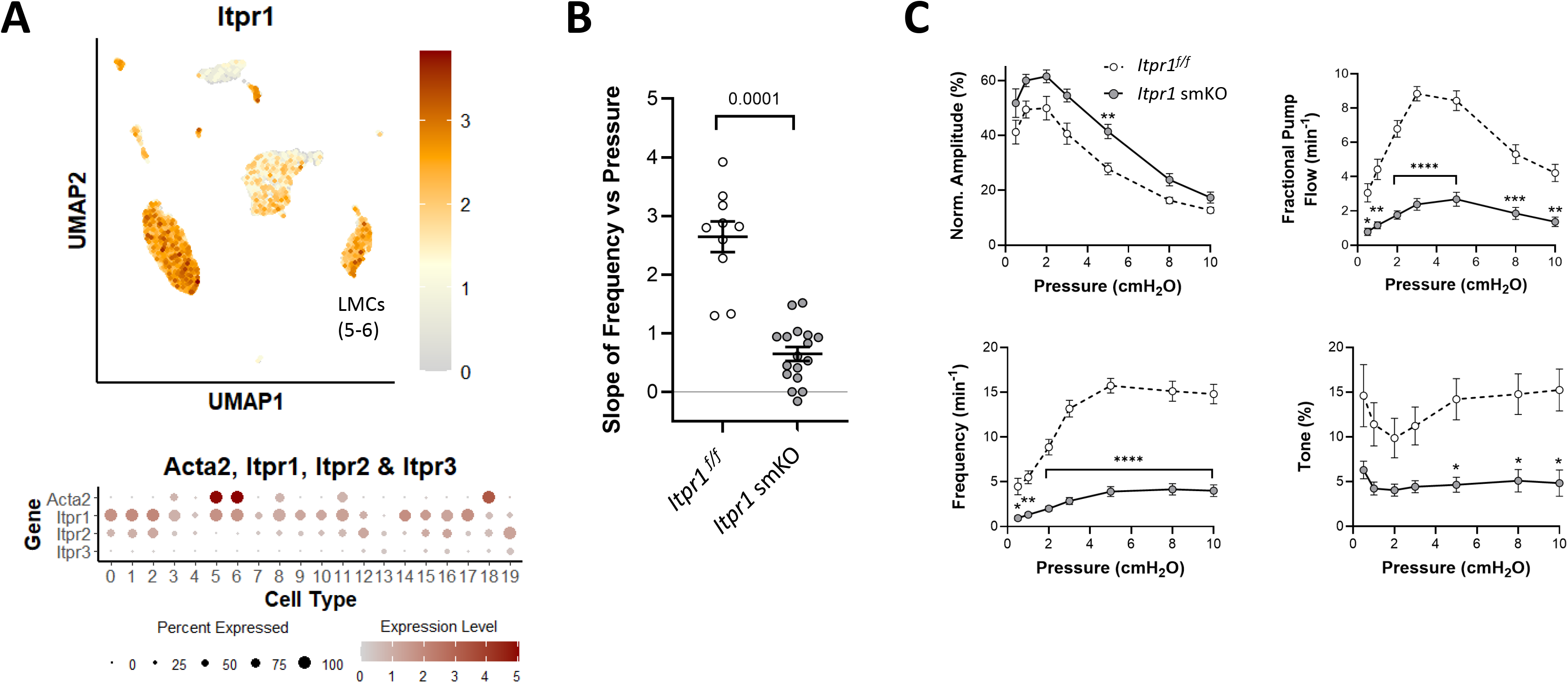
Consequences of *Itpr1* (IP_3_R1) deletion. UMAP and bubble plots of *Ip3r1* expression in the various IALV cell clusters. **B**) Slope of F-P relationship for popliteal lymphatics from *Itpr1^f/f^ and Itpr1* smKO mice. **C**) Summary plots of contraction parameters as a function of pressure for the two respective groups of vessels. Horizontal bar indicates range of pressures with same specified statistical significance. *Itpr1* smKO N=5, n=8; *Itpr1^f/f^* N=5, n=10.

**Figure 17.**
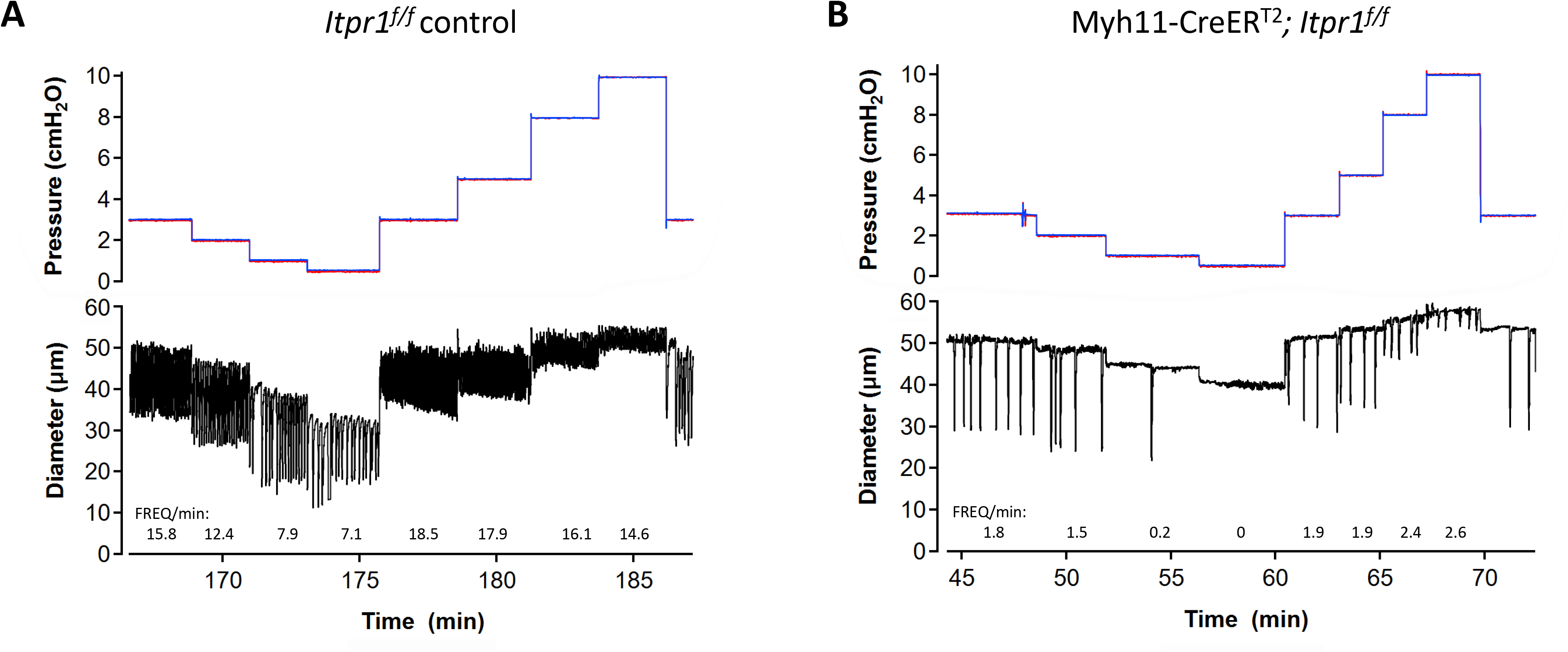
Contraction patterns of *Itpr1^f/f^* and *Itpr1* smKO vessels. A) Spontaneous contractions of an *Itpr1^f/f^* (control) popliteal lymphatic vessel at various pressures. Average frequencies at each pressure are listed below the diameter traces. Frequency ranged from 7.1 min^-1^ at 0.5 cmH_2_O to 17.9 min^-1^ at 5 cmH_2_O. B) Spontaneous contractions of an Myh11-CreER^T2^*;Itpr1^f/f^* (*IP_3_R1* smKO) popliteal lymphatic vessel at various pressures. Frequency ranged from 0 min^-1^ at 0.5 cmH_2_O to 1.9 min^-1^ at 5 cmH_2_O.

### Role of GNAQ/GNA11 in pressure-induced chronotropy

Evidence from multiple studies specifically points to GNAQ/GNA11-coupled GPCRs as the upstream mechanosensing elements of myogenic constriction in arteries (Mederos y Schnitzler *et al*., 2008; Blodow *et al*., 2014; Harraz *et al*., 2014; Li *et al*., 2014b; Schleifenbaum *et al*., 2014; Pires *et al*., 2017; Bjorling *et al*., 2018). Mechanosensitive activation of one or more GPCRs independent of ligand binding (Zou *et al*., 2004; Erdogmus *et al*., 2019) could account for pressure-induced increases in IP_3_ levels in LMCs. Our scRNAseq analysis revealed strong expression of *Gnaq* and *Gna11* in 50-75% of LMCs, LECs, and AdvCs as well as very strong expression of *Gnaq* in some immune cell clusters (**Fig. 18A**). We tested the effects of SM-specific *Gnaq* knock out (Myh11-CreER^T2^*;Gnaq^f/f^*), global *Gna11* knock out (*Gna11^-/-^*) and *Gnaq*/*Gna11* double knock out (Myh11-CreER^T2^*;Gnaq^f/f^;Gna11^-/-^*) on pressure-induced lymphatic chronotropy. There was a trend for *Gnaq* smKO vessels to have a reduced slope in the F-P relationship in comparison to *Gnaq^f/f^;Gna11^+/+^*control vessels, but it did not reach statistical significance (p=0.0819). In contrast, global deletion of *Gna11* significantly reduced the slope of the F-P relationship (p=0.0304) and the reduction was even more significant and pronounced (p<0.0001) in *Gnaq/Gna11* DKO vessels (**Fig. 18B**). However, even combined knock out of both proteins did not cause an attenuation of pressure-induced chronotropy that was comparable to that observed after SM-specific *Itpr1* or *Ano1* deletion, particularly at pressures ≥ 3 cmH_2_O (compare **Fig. 18B** to **Figs. 2B, 16B**). Contraction data for *Gnaq* smKO, *Gna11^-/-^* and *Gnaq/Gna11* DKO vessels, compared to tamoxifen-treated *Gnaq^f/f^;Gna11^+/+^*controls, are shown in the other panels of **Fig. 18C**. The full F-P relationships for each genotype showed significant reductions in frequency at a few pressures for *Gnaq* smKO vessels compared to *Gnaq^f/f^;Gna11^+/+^* controls, significant differences for *Gna11^-/-^* vessels at more pressures and significant differences at all pressures for *Gnaq*/*Gna11* DKO vessels. Collectively, these results suggest that both GNAQ and GNA11-coupled G-proteins are involved in pressure-induced lymphatic chronotropy, with G11 being more important. However, the observations that frequencies were higher than those in *Ano1* smKO and *Itpr1* smKO vessels (compare frequency plots in **Fig. 18C** to **Figs. 2C, 16C**) and that frequency continued to moderately increase at higher pressures when Gq/11 signaling is abrogated point to the possible contribution of other mechanisms. Examples of the lymphatic contraction patterns at each pressure in a *Gnaq^f/f^;Gna11^+/+^* control vessel and a *Gnaq*/*Gna11* DKO vessel are shown in **Figure 19**; in the DKO vessel the basal contraction frequency was only 1.1 min^-1^ at P=3 cmH_2_O and increased only to 1.3 min^-1^ at P=5 cmH_2_O.

**Figure 18.**
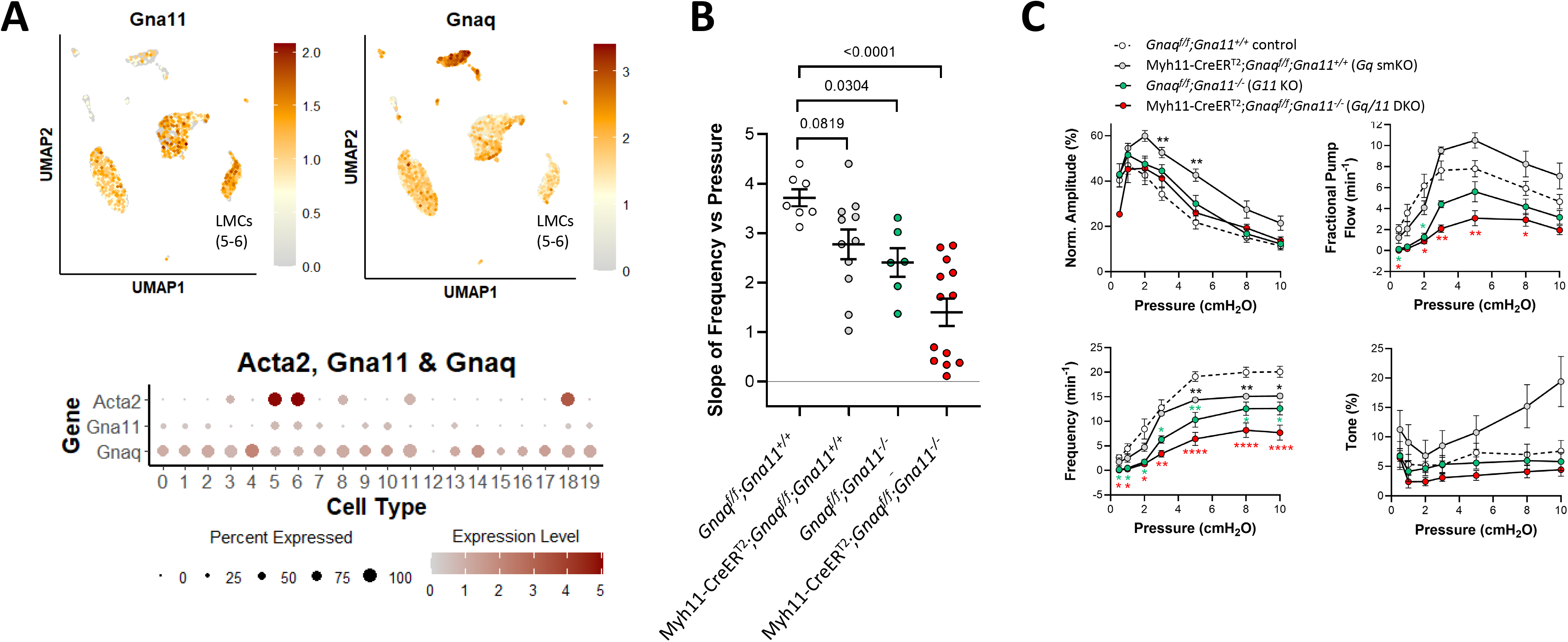
Consequences of *Gnaq* and/or *Gna11* deletion. UMAP and bubble plots of *Gnaq* and *Gna11* expression in the various IALV cell clusters. **B**) Slope of F-P relationship for popliteal lymphatics from *Gnaq^f/f^;Gna11^+/+^,* Myh11-CreER^T2^*;Gnaq^f/f^* (*Gnaq* smKO*), Gna11^-/-^ and* Myh11-CreER^T2^*;Gnaq^f/f^;Gna11^-/-^ (Gnaq/Gna11* DKO) mice. **C**) Summary plots of contraction parameters as a function of pressure for the four respective groups of vessels. *Gnaq^f/f^;Gna11^+/+^* control N=3, n=6; *Gnaq* smKO N=4, n=8; *Gna11^-/-^* N=3, n=6; Myh11-CreER^T2^*;Gnaq^f/f^;Gna11^-/-^*N=8, n=13.

**Figure 19.**
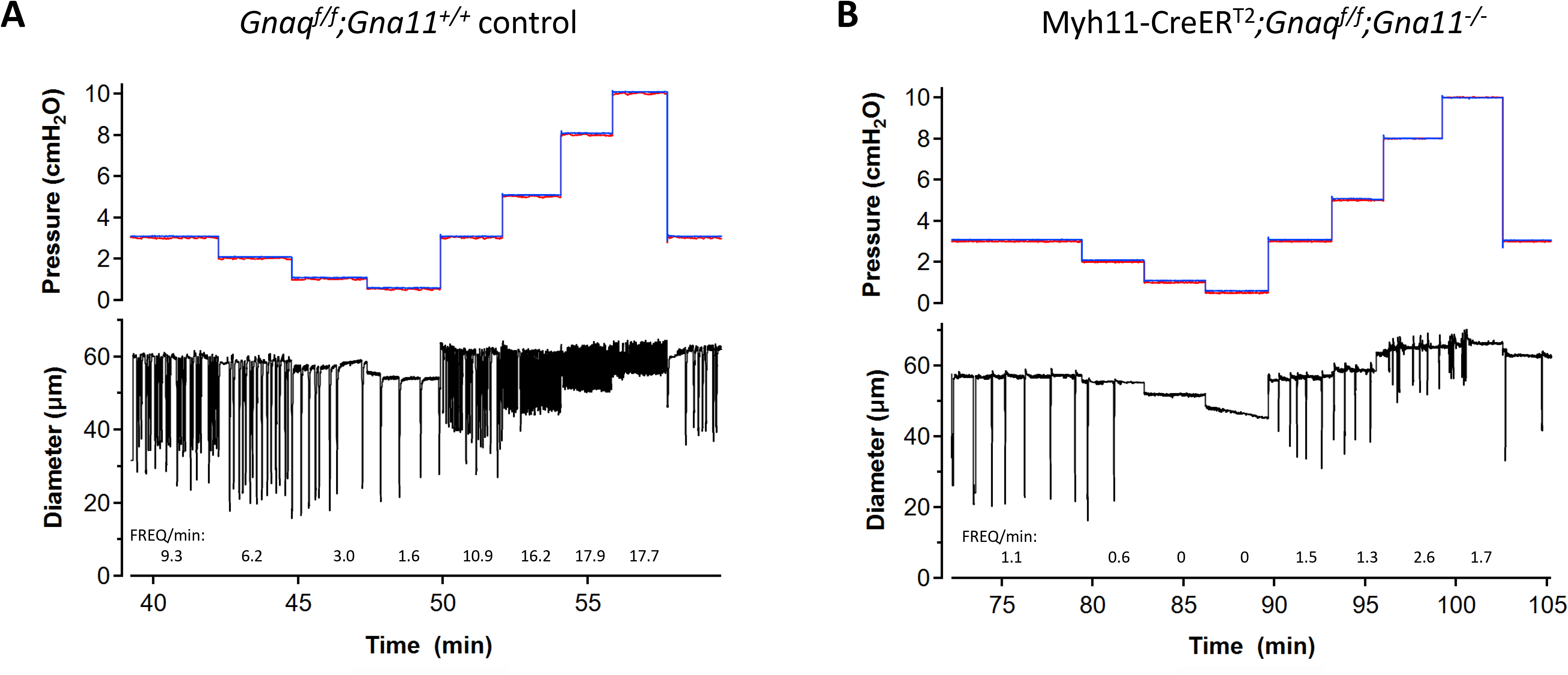
Contraction patterns of *Gnaq^f/f^;Gna11^+/+^* and Myh11-CreER^T2^*;Gnaq^f/f^;Gna11^-/-^*vessels. A) Spontaneous contractions of a *Gnaq^f/f^;Gna11^+/+^* (control) popliteal lymphatic vessel at various pressures. Average frequencies at each pressure are listed below the diameter traces. Frequency ranged from 1.6 min^-1^ at 0.5 cmH_2_O to 16.2 min^-1^ at 5 cmH_2_O. B) Spontaneous contractions in a Myh11-CreER^T2^*;Gnaq^f/f^;Gna11^-/-^* popliteal vessel at various pressures. Frequency ranged from 0 min^-1^ at 0.5 cmH_2_O to 1.3 min^-1^ at 5 cmH_2_O. The noisy diameter signals at P=8 and 10 cmH_2_O are caused by tracking glitches and do not represent high frequency contractions.

### Role for GNA12/GNA13 in pressure-induced chronotropy

GNA12/GNA13-coupled GPCRs are known to regulate arterial smooth muscle contraction through Rho/Rho kinase (Bolz *et al*., 2003; Dubroca *et al*., 2005; Gokina *et al*., 2005; Jarajapu & Knot, 2005; Li & Brayden, 2017; McDuffie *et al*., 2024) and at least one study suggests that Rho kinase is essential for phasic lymphatic contractions (Kurtz *et al*., 2014). scRNAseq analysis revealed weak expression of *Gna12* and *Gna13* in ∼25% of LMCs, with stronger expression of *Gna13* in LECs and AdvCs (**Fig. 20A**). Chennupati et al. (Chennupati *et al*., 2019) made the surprising finding that GNA12/GNA13-coupled GPCRs, rather than GNAQ/GNA11-coupled GPCRs, mediated myogenic constrictions in mesenteric and cerebral arteries of mice. We investigated the effects of deleting both *Gna12* and *Gna13* from LMCs (Myh11-Cre^T2^*;Gna12^f/f^;Gna13^-/-^*mice) on the F-P relationship of popliteal lymphatics. In the *Gna12*/*Gna13* double knock out, contraction amplitude was significantly blunted and tone was significantly elevated at several lower pressures, but the F-P relationship was nearly identical to that of vessels from control *Gna12^f/f^;Gna13^+/+^*mice (**Fig. 20B**) and frequencies of *Gna12*/*Gna13* DKO vessels were nearly identical to those of controls at all pressures (**Fig. 20C**). Thus, GNA12/GNA13-coupled GPCRs do not appear to be essential for pressure-induced chronotropy.

**Figure 20.**
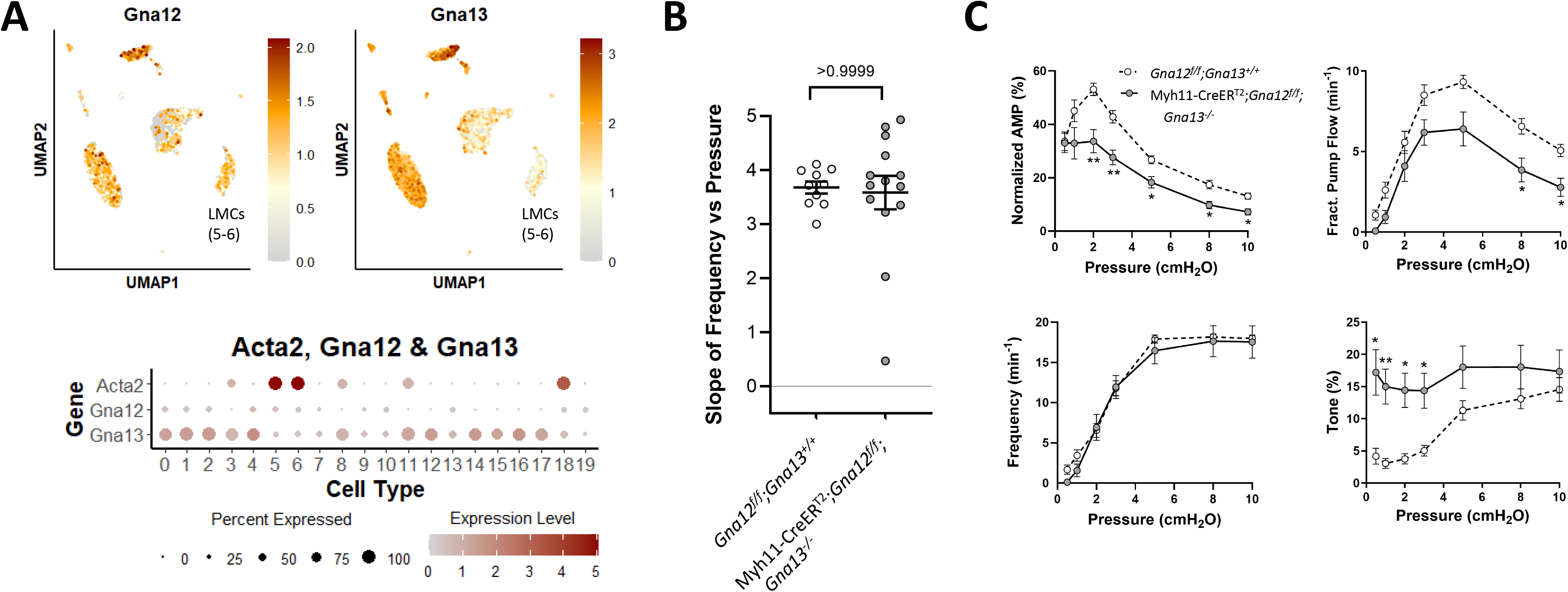
Consequences of *Gna12*/*Gna13* deletion. UMAP and bubble plots of *Gna12* and *Gna13* expression in the various IALV cell clusters. **B**) Slope of F-P relationship for popliteal lymphatics from *Gna12^f/f^;Gna13^+/+^ and* Myh11-CreER^T2^*;Gna12^f/f^;Gna13^-/-^*mice. **C**) Summary plots of contraction parameters as a function of pressure for the two respective groups of vessels. *Gna12^f/f^;Gna13^+/+^* controls N=5, n=10; Myh11-CreER^T2^*;Gna12^f/f^;Gna13^-/-^*N=7, N=15.

### Roles of specific GPCRs in pressure-induced chronotropy

The findings in **Fig. 18** strongly implicate Gq/11 signaling in pressure-induced lymphatic chronotropy but leave open the question of which specific GPCR(s) mediate(s) this mechanosensitive response. An scRNAseq analysis revealed that 136 GPCRs were expressed at various levels in mouse LMCs, using an expression threshold minimum of 0.5% LMCs. We constructed a bubble plot showing the top 20 of these GPCRs, ranked according to their percent expression across the LMC populations (**Fig. 21**, lines 11-30) and included GPCRs implicated in mechanosensation of the arterial myogenic response (32-41). The list includes the NPY1 receptor (*Npy1r*, line 11), the endothelin type A receptor (*Ednra*, line 13), the atrial natriuretic peptide type C receptor (*Npr3*, line 19), the thromboxane A2 receptor (*Tbxa2r*, line 20), the histamine type1 receptor (*Hrh1,* line 40) and the sphingosine-1-phosphate receptor 1 (*S1pr1*, line 38). Several of the more highly expressed GPCRs are members of the adhesion GPCR subfamily (*Adgrl1, Adgrl2, Adgre5, Adgrd1, Adgrl3, Adgra2; rows 15, 12, 17, 24-26*), and two more are orphan receptors (*Gprc5b, Olfr1033; rows 27,18*). Both subtypes of AT_1_R are expressed, but only at very low levels <1% (lines 32-33).

**Figure 21.**
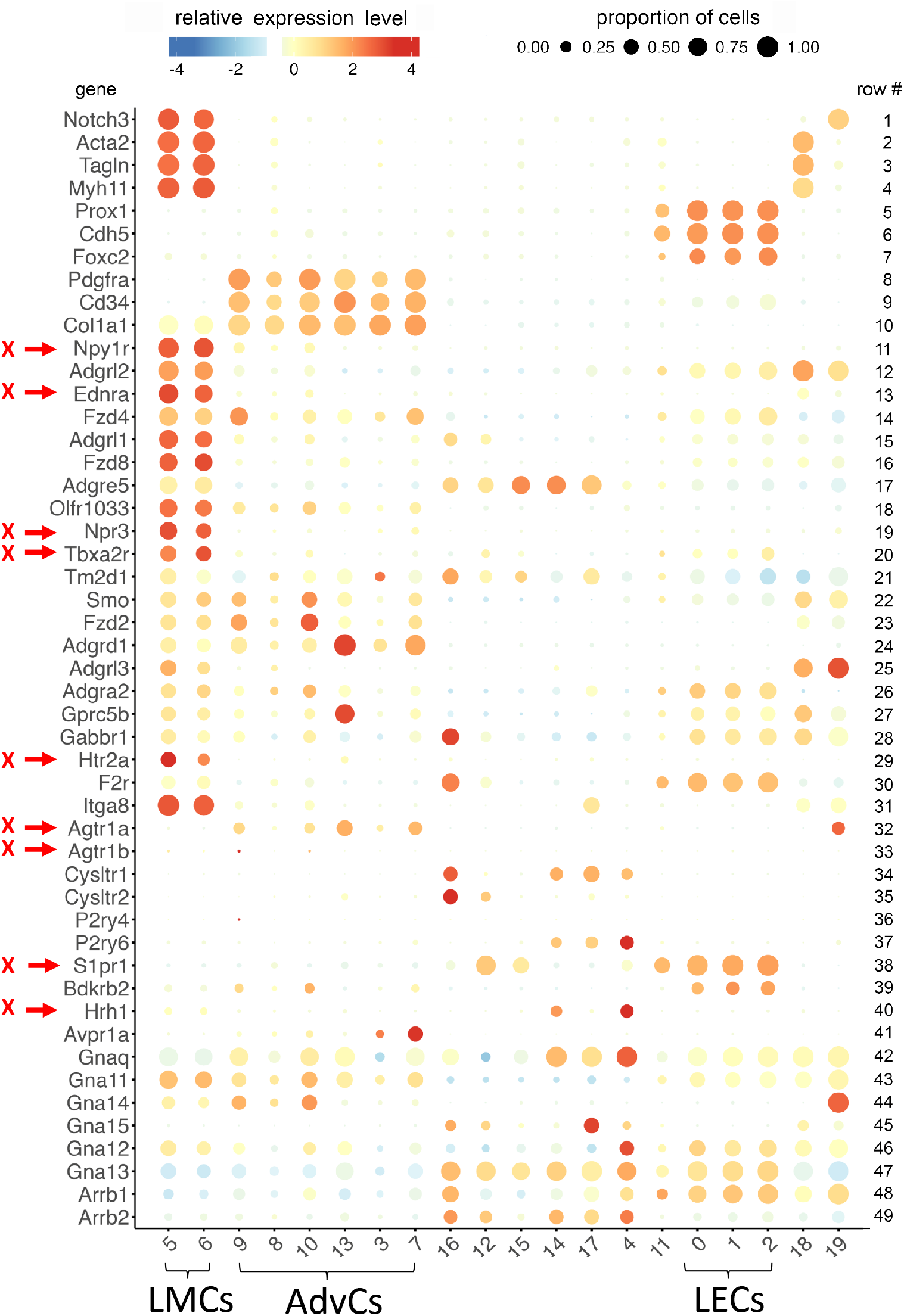
Bubble plot representing the results of scRNAseq analysis of the most highly expressed GPCRs in the various cell types. Rows 1-10 show the canonical markers for LMCs (rows 1-4), LECs (rows 5-7) and AdvCs (rows 8-10), respectively. Rows 11-30 show the top 20 GPCRs expressed in LMCs, ranging from 87.5% of LMCs expressing *Npy1R* to 36.6% LMCs expressing *F2R*. Putative mechanosensitive GPCRs in other cell types (e.g. arterial SM), but with very low or absent expression in LMCs, are shown in rows 34-41; AT_1_R subtypes are shown in rows 32-33, cysteinyl leukotriene receptor subtypes in rows 34-35 and P2Y receptor subtypes in rows 36-37. Four other GPCRs are shown in rows 38-41. G-coupling proteins are shown in rows 42-47 and β-arrestins in rows 48-49. Red arrows at left are the GPCRs that were tested with inhibitors in the present or previous studies (see text for references).

We proceeded to experimentally test the involvement of several of the above GPCRs for which suitable receptor antagonists were available. In lieu of evaluating the complete pressure-frequency response for each GPCR/antagonist, we determined whether blocking the respective GPCR would alter pacemaking by lowering the contraction frequency at P=3 cmH_2_O. As confirmation that this procedure was sufficiently sensitive to detect an impact on normal pacemaking frequency, we compared the frequencies at 3 cmH_2_O for the *Ano1* smKO, *Itpr1* smKO and *Gnaq/Gna11* DKO vessels and their respective controls (**Fig. 22**, gray symbols). We also performed this analysis for WT vs Ani9-treated vessels and for *Gnaq^+/+^;Gna11^+/+^* control vessels before and after treatment with the highly selective and potent Gq/11 antagonist YM254890 (Takano *et al*., 2025). As expected, each of those comparisons was highly significant, indicating that the inhibitor or knock out reduced contraction frequency. Likewise, YM254890 significantly reduced the frequency of *Gnaq^+/+^;Gna11^+/+^*vessels, providing further confirmation that Gq/11 signaling is critically important for the chronotropic response. However, *none* of the GPCR antagonists, including those for the ANP receptor, the ET1_A_ receptor, the NPY1 receptor, the thromboxane A2 receptor, or S1P type 1 receptor, produced a significant reduction in frequency (**Fig. 22**, red symbols). We also compared the responses of vessels from *Agtr1a^-/-^* mice and from WT mice treated with losartan, which blocks both AT_1a_R and AT_1b_R; in both the KO and the WT + losartan vessels there was actually a significant *increase* in contraction frequency. We conclude that none of these seven GPCRs are critical for pressure-induced chronotropy of mouse collecting lymphatic vessels.

**Figure 22.**
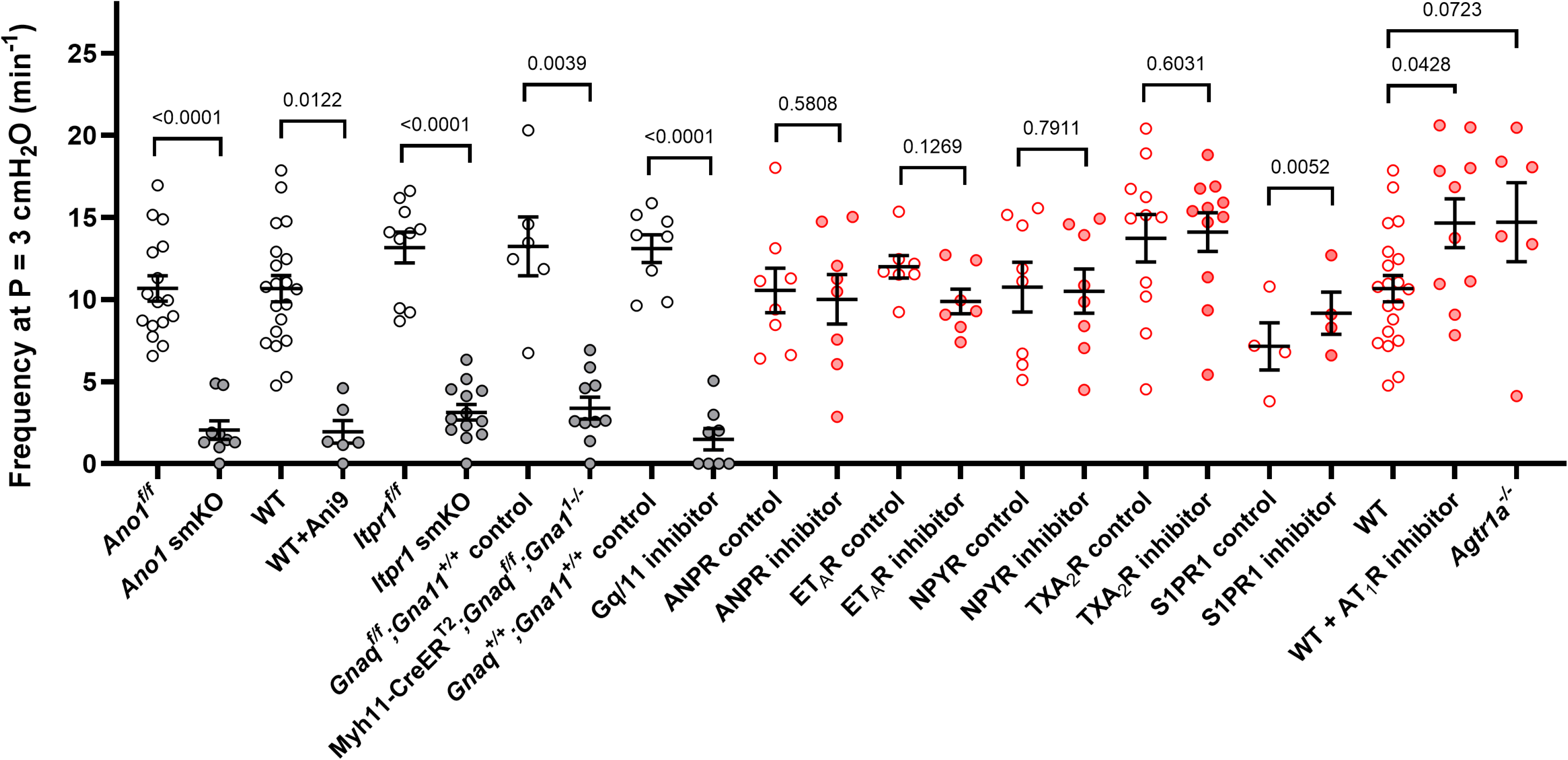
Screening the effects of selective GPCR inhibitors. Pacemaking frequency is compared between control and inhibitor (or knock out) at a single pressure, 3 cmH_2_O. Gray symbols: comparisons between *Ano1^f/f^* controls and *Ano1* smKO, WT ± Ani9, *Itpr1^f/f^* controls and *Itpr1* smKO, *Gnaq^f/f^;Gna11^+/+^* controls and Myh11-CreER^T2^*;Gna12^f/f^;Gna13^-/-^* vessels, *Gnaq^+/+^;Gna11^+/+^* controls ± YM254890, showing that each treatment produced a very significant reduction in frequency using this test. Red symbols: comparisons between WT controls ± respective inhibitors for the various GPCR. None of the GPCR inhibitors produced a significant decrease in frequency, but the S1PR1 and AT_1_R inhibitors instead significantly increased frequency. Inhibitor concentrations were as follows: Gq/11 inhibitor YM254890 (100 nM); ANPR inhibitor AP811 (1 μM); ET1_A_R inhibitor BQ123 (1 μM); NPYR inhibitor BIBO3304 (1 μM); TXA_2_R inhibitor S18886 (20 nM); S1PR1 inhibitor SEW-2871 (1 μM); AT_1_R inhibitor losartan (10 μM). Comparisons between controls and KO vessels were made using 2-way ANOVAs with Dunnett’s post-hoc tests. Comparisons between controls ± inhibitors were made using paired t-tests. *Ano1* smKO N=14, n=23; *Ano1^f/f^* N=8, n=16. WT N=15, n=20; WT ± Ani9 N=5, n=6. Myh11-CreER^T2^*;Itpr1^f/^*N=5, n=8; *Itpr1^f/f^* N=5, n=10. *Gnaq^f/f^;Gna11^+/+^* controls N=3, n=6; Myh11-CreER^T2^*;Gnaq^f/f^;Gna11^-/-^* N=8, n=13. *Gnaq^+/+^;*Gna11^+/+^ controls ± YM254890 N=8, n=8. WT ± AP811 N=8, n=8. WT ± BQ123 N=7, n=7. WT ± BIBO3304 N=8, n=8. WT ± SEW-2871 N=4, n=4. WT ± losartan N=5, n=10. *Agtr1a^-/-^*N=3, n=6. S18886 N=4; n=11.

## Discussion

### Lack of role for mechanosensitive ion channels in pressure-induced lymphatic chronotropy

The major goal of this study was to investigate the mechanistic basis of pressure-induced lymphatic chronotropy. Because little was previously known regarding the mechanisms of mechanosensitivity underlying this phenomenon, we took cues from a well-studied mechanosensitive mechanism in vascular smooth muscle—the arterial myogenic response. In both arteries and collecting lymphatic vessels (Davis *et al*., 2009; Davis *et al*., 2023d), pressure elevation leads to vascular muscle cell depolarization and constriction, reflecting the activation of a depolarizing current that triggers Ca^2+^ entry through the opening of L-type, voltage-gated Ca^2+^ channels. Several ionic mechanisms have been previously implicated in this process for arterial smooth muscle, specifically the activation of TRPC6, TRPM4, PKD1/2, TRPV2 and ENaC cation channels (Welsh *et al*., 2002; Earley *et al*., 2004a; Jernigan & Drummond, 2005; Sharif-Naeini *et al*., 2009; McGahon *et al*., 2016; Bulley *et al*., 2018), with supporting evidence for each obtained from gene knockdown/knockout studies. Additional ion channels might also be involved, including the activation of channels promoting Cl^-^ efflux (Boedtkjer *et al*., 2008) or the inhibition of constitutively active K^+^ channels (Sancho *et al*., 2019), but evidence for those is based largely on the use of pharmacological inhibitors and therefore subject to concerns about off-target effects (Maroto *et al*., 2005; Dietrich *et al*., 2007; Gottlieb *et al*., 2008; Sancho *et al*., 2019). In the present study, we primarily used genetic knock out approaches to test the possible roles of the major ion channels contributing to the mechanosensitive chronotropy of lymphatic vessels from the mouse. Our results strongly support the conclusion that pressure-induced lymphatic chronotropy does *not* involve acute activation of any of the putative mechanosensitive (and/or second-messenger gated) cation channels previously implicated in myogenic constriction (TRPC6, TRPM4, PKD2, TRPV2, ENaC), nor does it require PIEZO1, the bona fide mechanosensitive cation channel implicated in other aspects of mechanotransduction in the cardiovascular system [see references in (Davis *et al*., 2023d)]. Instead, our findings support a critical role for calcium dynamics regulating ANO1 in pressure-induced lymphatic chronotropy. ANO1 is a channel that is also required for the generation of arterial myogenic tone (Bulley *et al*., 2012). Studies of mammalian ANO1 indicate that calcium store release is the primary mode of ANO1 regulation (Xiao *et al*., 2011; Dulin, 2020) and our observation that IP_3_R1 knock out has an almost identical effect on pressure-induced chronotropy as ANO1 deletion supports the conclusion that IP_3_-induced calcium release from SR stores through IP_3_R1 drives ANO1 activation in LMCs (Zawieja *et al*., 2023). Our further observation that Gq/11 deletion or inhibition produces nearly equivalent effects on pressure-induced chronotropy as ANO1 or IP_3_R1 knock out—especially at lower pressures—are consistent with pressure-induced phospholipase activation and IP_3_ production (Narayanan *et al*., 1994) that is driven by an upstream mechanosensor. Based on an extensive body of literature in cardiomyocytes (Yasuda *et al*., 2008) and arterial smooth muscle (Schleifenbaum *et al*., 2014; Cui *et al*., 2022), mechanotransduction of stretch/pressure is mediated by one or more GNAQ/GNA11-coupled GPCRs, leading to downstream activation of second messenger-gated channels. In the case of LMCs, mechano-activation of GNAQ/GNA11-coupled GPCRs would promote IP_3_ production that triggers Ca^2+^ release from stores through IP_3_R1; Ca^2+^ would then activate ANO1 Cl^-^ channels, which would accelerate the rate of spontaneous diastolic depolarization, increasing the firing rate of Cav1.2 action potentials that control phasic lymphatic contractions. Repolarization is known to be controlled by Kv channels (Cotton *et al*., 1997), including Kv11 channels (Kim *et al*., 2023). As summarized in **Table 1**, knockout of each element in the Gq/11-IP_3_-IP_3_R1/Ca^2+^-ANO1-Cav1.2 pathway (and only those elements out of all that were tested, except *Kcnj2/ Kcnj12*, which can be alternatively explained) abrogated or significantly attenuated pressure-induced chronotropy.

**Table 1.** Genes for which deletion/inhibition produced significant attenuation or enhancement of the F-P relationship.

| **Attenuation** | **Enhancement** |
|---|---|
| *Cacna1c* | *Trpc6/Trpc3* |
| *Ano1* | - |
| *Itpr1* | - |
| *Gnaq* | - |
| *Gna11* | - |
| *Kcnj2/Kcnj12* | - |

Although we tested the requirement for 16 potential mechanosensitive ion channels in pressure-induced lymphatic chronotropy, our list was not exhaustive. Many channels have some degree of intrinsic mechanosensitivity [for review see (Davis *et al*., 2023d)] but mechanosensitivity for most has been demonstrated only under quite artificial conditions (e.g., patch clamp + localized membrane suction or hypoosmotic swelling) in which the mechanical stimulus is unlikely to accurately mimic the magnitude and specificity of force experienced by that channel in a cell in vivo (Davis *et al*., 2023d). The list of bona fide mechanosensitive channels is very short if one adheres to the criteria proposed by Patapoutian and colleagues (Ranade *et al*., 2015). Cell volume changes can activate other classes of ion channels, but the volume of an LMC is unlikely to substantially change during the small pressure steps that produce increased chronotropy. Nevertheless, we also tested the potential roles of selected volume activated channels implicated in arterial myogenic constriction, including TRPV2, TRPV4 and Kir (McGahon *et al*., 2016; Soni *et al*., 2017; Sancho *et al*., 2019), but not others reported to be important in other cell types or in other physiological contexts (Matchkov *et al*., 2015). Another channel known to influence LMC pacemaking that was not explicitly tested here is the voltage-gated sodium channel (VGNaC), Nav1.3.

VGNaCs are present and active in LMCs, with Nav1.3 most consistently being expressed (Telinius *et al*., 2015; Zawieja *et al*., 2024; Arroyo-Ataz *et al*., 2025; Ruscic *et al*., 2025). VGNaC contributions to pacemaking appear to vary by species, in part because of differences in the resting membrane potential (Davis & Zawieja, 2024). Telinius et al. found that APs and spontaneous contractions were inhibited by the selective VGNaC blocker, TTX, in 60% of human mesenteric lymphatics (Telinius *et al*., 2015). In contrast, phasic contractions of guinea pig and sheep mesenteric lymphatics continued in the presence of TTX (Chan & von der Weid, 2003; Chan *et al*., 2004; Beckett *et al*., 2007). We previously found that TTX had no detectable effect on phasic contractions in ∼50% of mouse popliteal lymphatic vessels and only a transient inhibitory effect (lasting 1-4 minutes) in the continued presence of the toxin in the other ∼50% (Davis *et al*., 2023c). In mouse IALVs, a recent study reported similar, non-detectable effects of TTX (Schulz *et al*., 2026) and Padera and colleagues report that spontaneous lymphatic pumping persists in popliteal lymphatics of *Nav1.3* KO mice (Ruscic *et al*., 2025). Thus, Nav1.3 channels do not appear to be critical for pressure-induced chronotropy of mouse lymphatic vessels.

### Roles for TRP channels in other aspects of LMC contractile function

If IP_3_ production in LMCs is enhanced after pressure elevation, as in arterial smooth muscle (Narayanan *et al*., 1994; Mederos y Schnitzler *et al*., 2008), then concomitant increases in DAG, the other product of PIP_2_ hydrolysis, would also be predicted to occur and DAG should then activate TRPC6 and related channels. However, if this mechanism is operating in popliteal lymphatic muscle, the activation of those channels apparently does not significantly influence the F-P relationship (**Fig. 4**). Likewise, the Ca^2+^ increase resulting from mechanosensitive activation of the Gq/11-IP_3_-IP_3_R1 signaling axis would potentially activate TRPM4 and/or PKD2 channels (assuming they are localized to the appropriate LMC microdomains), but we find that their deletion does not significantly alter the F-P relationship (**Figs. 5-6**). Because TRPM4, PKD2 and PIEZO1 expression levels are very low in LMCs (**Figs. 3A,4A**, **5A**), further activation of those channels may have little effect. It is possible that another unidentified ion channel could be mediating the pressure-driven frequency changes at higher pressures—perhaps a channel with a higher mechanosensitive threshold or requiring higher levels of IP_3,_ DAG or Ca^2+^. Candidates include TRPV1, which has also been implicated in arterial myogenic constriction (Phan *et al*., 2022).

Although the deletion/inhibition of several TRP channel family members and other channels did not significantly impact pressure-induced chronotropy, there was sometimes an impact on other aspects of lymphatic contractile function, namely contraction amplitude and/or tone. The respective channels and their statistically significant contraction effects are summarized in **Table 2**. Deletion of *Trpc6/Trpc3* impaired contraction amplitude by ∼40% and increased tone by ∼2.5-fold (**Fig. 4C**). Deletion of *Trpv4* from LMCs impaired contraction amplitude by ∼45% without an effect on tone (**Fig. 7C**). TRPV4 channel activation has recently been implicated in the regulation of lymphatic contraction, but primarily via the production of vasoactive products from LECs and resident macrophages (Schulz *et al*., 2025). Inhibitors of three other ion channels also led to significant changes in amplitude and/or tone. Specifically, one of the two TRPV2 inhibitors (SET2) inhibited tone at 4 of the 5 lowest pressures. One of the two ENaC inhibitors (amiloride) inhibited amplitude by 20-30% while the other (benzamil) enhanced tone by ∼2-fold (**Fig. 9C**). The Kir2 blocker Ba^2+^ impaired amplitude by ∼40% and increased tone by 2-fold (**Fig. 13C**). In the latter case the effects of Ba^2+^ on tone and frequency are most likely explained by the inhibition of endothelial Kir channels, which control the production of inhibitory vasoactive products (Scallan & Davis, 2013; Davis *et al*., 2023d; DuToit *et al*., 2024; Davis & Bertram, 2025). Collectively, these observations suggest that the respective channels are each involved in some aspect of lymphatic contractile function other than pressure-induced chronotropy, with the more compelling evidence being obtained after gene deletion rather than after acute application of soluble inhibitors, which may have off-target effects.

**Table 2.** Consequences of deletion/inhibition of ion channels on contraction amplitude and/or tone.*

| **Channel** | **Effect on AMP** | **Effect on Tone** | **Method of inhibition** |
|---|---|---|---|
| TRPC6/3 | ↓ | ↓ | deletion |
| TRPV4 | ↓ | - | deletion |
| TRPV2 | - | ↓ | soluble inhibitor |
| ENaC | ↓ | ↑ | soluble inhibitor |
| Kir2.x | ↓ | ↑ | soluble inhibitor |

\* Excluding the GNAQ/GNA11-IP<sub>3</sub>R1/Ca<sup>2+</sup>-ANO1-Cav1.2 signaling pathway.

### Pressure-induced lymphatic chronotropy requires intact Gq/11 signaling

Our results using mice with SM-specific knock out of *Gnaq* and/or global knock out of *Gna11* suggest that both Gq and G11 contribute to pressure-induced chronotropy. The F-P slope of popliteal lymphatics was significantly reduced in *Gna11^-/-^* vessels and more so in *Gnaq/Gna11* DKO vessels (**Fig. 18**), as well as in WT vessels treated with the specific Gq/G11 inhibitor YM254890 (**Fig. 22**). Although the reduction in F-P slope for *Gnaq* smKO vessels did not reach statistical significance (**Fig. 18A**), contraction frequencies at 3 of 7 pressures were significantly lower than in *Gnaq^f/f^* controls (**Fig. 18C**). Additionally, the F-P slope for *Gq/11* DKO vessels was lower than that of *Gna11^-/-^* vessels, so it appears that the effects of knocking out *Gnaq* and *Gna11* are cumulative, suggesting that *Gnaq* deletion contributes to the lower F-P slope of *Gnaq/Gna11* DKO vessels. Further support for this conclusion is provided by a similar analysis of IALVs in which *Gnaq/Gna11* DKO vessels had significantly lower frequencies than *Gna11^-/-^* vessels at 3 of the 4 lowest pressures (**Fig. 18**. **Suppl. 1**). Further inspection of **Figure 18C** suggests that frequency is impaired at high pressures in *Gnaq* smKO popliteal vessels whereas frequency is impaired at low pressures in *Gna11^-/-^* popliteal vessels, so the two G-coupled proteins may associate with GPCRs that sense different pressure ranges.

Our conclusions are consistent with evidence that Gq/11-coupled GPCRs are required for mechanotransduction of the arterial myogenic response. Pressure-induced constriction in several types of arteries is significantly attenuated or even completely blocked by inverse agonists of AT_1_R (Mederos y Schnitzler *et al*., 2008; Schleifenbaum *et al*., 2014; Storch *et al*., 2015; Hong *et al*., 2016) and/or by knock-out of the specific receptor subtypes AT_1A_R (Schleifenbaum *et al*., 2014; Cui *et al*., 2022) or AT1_B_R (Pires *et al*., 2017). However, although AT_1_R may be a critical mechanosensitive GPCR in many arteries, other studies have implicated additional GPCRs, including the cysteinyl leukotriene 1 receptor (Storch *et al*., 2015; Mederos *et al*., 2016) and P2Y4/P2Y6 receptors (Brayden *et al*., 2013; Forst *et al*., 2016; Kauffenstein *et al*., 2016). None of those GPCRs are likely candidates to mediate pressure-induced lymphatic chronotropy as they are all expressed at extremely low levels in only a small fraction of LMCs (**Fig. 21**, lines 34-37). Likewise, knock out of *Agtr1a* did not significantly alter the lymphatic vessel F-P relationship, nor did the AT_1_R antagonist losartan, which inhibits both AT_1A_R and AT_1B_R subtypes (**Fig. 22**), indicating that AT_1_R is not required for this aspect of lymphatic mechanotransduction. Other Gq/11-coupled GPCRs in VSMCs that are potentially mechanosensitive include S1PR1, B2R, H1R, M5R and V1AR (Mederos y Schnitzler *et al*., 2008; Peter *et al*., 2008). However, B2R, M5R and V1AR are expressed only at very low levels in LMCs (**Fig. 21**). We tested the S1PR1 inhibitor, SEW 2871, which did not inhibit contraction frequency but instead increased it (**Fig. 22**). Inhibition of H1R in two other studies failed to inhibit spontaneous lymphatic contractions (Fox & von der Weid, 2002; Nizamutdinova *et al*., 2014).

In arteries, blockade of Gi or Gs signaling fails to inhibit mechanosensitive VSM responses (Mederos y Schnitzler *et al*., 2008). However, an exception to Gq/11-mediated mechanotransduction in arterial smooth muscle is a study by Chennupati et al. (Chennupati *et al*., 2019) who found that G12/13 but not Gq/11 signaling was critical for myogenic responses of mouse mesenteric and cerebral arteries. To our knowledge, the conflict between the latter result and the body of literature supporting a Gq/11 mechanism (Mederos *et al*., 2016), even with respect to specific types of arteries, has not been adequately resolved. A requirement for G12/13 signaling fits with multiple studies suggesting a critical role for Rho/Rho kinase (downstream from G12/13) in arterial myogenic constriction (Bolz *et al*., 2003; Dubroca *et al*., 2005; Gokina *et al*., 2005; Jarajapu & Knot, 2005; Li & Brayden, 2017) and even for phasic lymphatic contractions (Kurtz *et al*., 2014). However, our finding that *Gna12/Gna13* DKO vessels exhibit normal pressure-induced lymphatic chronotropy adds further support for a Gq/11-coupled mechanism rather than a G12/13-coupled mechanism. We cannot rule out a role for G14 participating in Gq/11-associated GPCRs as *Gna14* is expressed in 28% of LMCs in our dataset (**Fig. 21**) and we did not test the effects of *Gna14* deletion. Further experiments testing whether each G-coupled protein is activated at differing pressure thresholds of the pressure range, or how gene dosage of *Gnaq/Gna11/Gna14* regulates pressure dependent chronotropy are required. Lastly, GPCRs are known to couple promiscuously, so there is likely crosstalk and overlap between mechanosensitive GPCRs and their coupling proteins in any given tissue (Hauser *et al*., 2022).

We were unable to identify a specific GPCR in lymphatic muscle that mediates IP_3_/Ca^2+^-activation of ANO1 in response to pressure elevation. Our scRNAseq analysis of the GPCRs expressed in mouse LMCs identified 136 GPCRs, and we listed the top 20 by percent expression in LMCs (rows 11-32 in **Fig. 21**). We then experimentally tested 7 of the most likely candidates for which inhibitors were available by determining whether blocking the GPCR would alter pacemaking by lowering the contraction frequency at P=3 cmH_2_O; the results for each of those GPCRs were negative (**Fig. 22**). Two additional GPCRs (HT-2R, HRH1) can be ruled out by previous experiments showing that pacemaking persisted in the presence of their respective receptor blockers (Chan & von der Weid, 2003; Nizamutdinova *et al*., 2017). We conclude that *none* of these nine GPCRs (NPY1R, ET_A_R, NPR3, TBAX2R, HT-2R, HRH1, S1PR1, AT1_A_R or AT1_B_R), are critical for pressure-induced chronotropy in LMCs (**Fig. 21**, red arrows at left). Of the remaining GPCRs expressed at modest to high levels in mouse LMCs (**Fig. 21**), several are adhesion receptors *Adgrl1, Adgrl2, Adgre5, Adgrd1, Adgrl3, Adgra2*), two are orphan receptors (*Gprc5b, Olfr1033*), one codes for the GABA-B1 receptor (*Gabbr1*, with unknown function in the lymphatic system), two code for proteins in the Wnt signaling pathway (*Fzd4, Fzd2*), one codes for a protein in the Hedgehog signaling pathway (*Smo*), and one encodes a beta-amyloid binding protein (*Tm2d1*). The possible roles of these GPCRs, and the other 100+ GPCRs expressed at much lower levels remain to be investigated in future studies, but in many cases testing will be limited by the lack of known ligands/antagonists. We also cannot rule out a GPCR-independent mechanism involving Gq/11-coupled proteins (Dela Paz *et al*., 2017), although the evidence for such a mechanism is weak (Cui *et al*., 2022).

### Lack of role for the endothelium and endothelial cell GPCRs in pressure-induced chronotropy

Is there a possible role for mechanotransduction of pressure by GPCRs in LECs? Some early vascular studies suggested that pressure-induced arterial constriction was mediated by factors released from the endothelium (Harder, 1987; Harder *et al*., 1989; Harder *et al*., 1990), however subsequent work in multiple laboratories was unable to confirm this as a general mechanism (Kuo *et al*., 1990; Davis *et al*., 2023d). Perhaps the best evidence for mechanosensitive GPCRs comes from studies of the endothelium in which multiple Gq/11-coupled GPCRs are implicated in shear-stress dependent BEC calcium signaling (Chachisvilis *et al*., 2006; Xu *et al*., 2018; Erdogmus *et al*., 2019), which drives the subsequent production of vasoactive products (Davis *et al*., 2023d). Elegant experiments with cultured BECs expressing truncated or modified GPCRs, including H1R, A2AR or BDKRB2, point to Helix 8 in the GPCR C-terminus as the common mechanosensitive region (Erdogmus *et al*., 2019). GPCRs without Helix 8 are not mechanosensitive and transfer of Helix 8 to non-mechanosensitive GPCRs endows mechanosensitivity (Erdogmus *et al*., 2019). As in VSMCs, Gi and Gs-coupled GPCRs appear not to be involved in BEC mechanotransduction (Gudi *et al*., 1996; Gudi *et al*., 1998). Because several of the Gq/11 coupled GPCRs in BECs listed above are also expressed in mouse LECs (**Fig. 21**), is it possible that LECs are mediating pressure-induced chronotropy? Indeed, oscillatory shear stress is a known driver of several LEC signaling processes requiring GPCRs, including the adrenomedullin receptor, *Calcrl*, S1PR1 and Plexin D (Fritz-Six *et al*., 2008; Geng *et al*., 2025; Pang *et al*., 2026) and some laminar shear stress responses in blood endothelium are mediated by GPCRs, including PECAM-1, VE-Cadherin, VEGFR2 and Plexin D (Tzima *et al*., 2005; Mehta *et al*., 2020). However, all those responses involve longer term processes such as cell reorientation, NF-κB activation and valve development, rather than a rapid change in contraction frequency following a pressure change.

A possible role for these GPCRs could be ruled out by removing the lymphatic endothelium. We were unable to test whether an intact lymphatic endothelium is required for pressure-induced chronotropy in mouse popliteal lymphatic vessels because it is nearly impossible to selectively remove the endothelial layer from such small vessels containing one-way valves without damaging the overlying single layer of LMCs. However, studies using larger lymphatic vessels in other species or regions have previously demonstrated that lymphatic pacemaking and/or stretch-induced chronotropy remain intact after partial/complete removal or disruption of the endothelium using an air bolus (Fox & von der Weid, 2002; Ferrusi *et al*., 2004; DuToit *et al*., 2024) or thin wire (Zhang *et al*., 2007; von der Weid *et al*., 2014).

Additionally, pulsatile shear stress (which is the only type of shear stress present under the conditions of our in vitro experiments) is *inhibitory* rather than excitatory to both the frequency of lymphatic contractions and tone (Gasheva *et al*., 2006; Davis & Bertram, 2025)—opposite the response needed to explain pressure-induced chronotropy. We also found that LEC-specific deletion of *Piezo1*, the mechanosensitive ion channel mediating shear stress-induced calcium entry in arterial endothelium (Li *et al*., 2014a; Rode *et al*., 2017), did not significantly impair pressure-induced lymphatic chronotropy (Fig. 4). Additionally, activation of PIEZO1 with Yoda resulted in a nitric oxide-dependent inhibition of IALV contractions (Choi *et al*., 2024). LMC pacemaking is not impaired by endothelium-specific deletion of *Trpv4* (Schulz *et al*., 2025), a Ca^2+^-activated, Ca^2+^-permeable channel providing additional calcium entry after PIEZO1 activation (Swain & Liddle, 2021; Davis *et al*., 2023d). Finally, *Ano1*, the key effector of pressure-induced chronotropy, is not expressed in LECs (**Fig. 2A**).

### Residual pacemaking and partial rescue of the F-P relationship at high pressures

Deletion/blockade of ANO1, IP_3_R1 or Gq/11 in LMCs was associated with the complete abolition of pacemaking and phasic contractions at low pressures (0.5-1 cmH_2_O) in ∼50% of popliteal lymphatic vessels (**Fig. 16**, **18**; F-P slope = 0). In other vessels, the pacemaking rate dropped to a low level that was nearly independent of pressure (**Fig. 2**; F-P slope <0.5) while in others, pacemaking frequency dropped to a low level but still increased to a fraction of its normal level when pressure was elevated to 8 or 10 cmH_2_O (F-P slope = 0.5-2). The residual pacemaking observed after removal of ANO1 activity likely represents the constitutive interactions between depolarizing current contributed by L-type Ca^2+^ channels and hyperpolarizing current contributed by K^+^ channels, as elucidated in recent models of lymphatic pacemaking (Hancock *et al*., 2022, 2023). These models describe coupled oscillations in the membrane voltage (the M-clock) and intracellular calcium concentration (the C-clock) in which both clocks are able to oscillate independently but with the M-clock normally driving the C-clock. Action potentials are thus triggered at a relatively constant basal rate but are subject to irregularities in pacing frequency due to small fluctuations in membrane potential and/or intracellular calcium release. Input from the Gq/11-IP_3_-IP_3_R1-Ca^2+^-ANO1 signaling axis results in a strong depolarizing input capable of accelerating the rate of oscillations (Hancock *et al*., 2022, 2023). The schematic in **Figure 23** summarizes this proposed mechanism in light of our findings in the present study. The ability of some vessels to respond to pressure elevation after ANO1, IP_3_R1, or Gq/11 blockade indicates that a Gq/11- and/or ANO1-independent mechanism is operating, or compensating, at higher pressures. However, pressures > 8 cmH_2_O are levels unlikely to be chronically experienced by mouse popliteal vessels (Davis, 2026) except perhaps in lymphedema. A potential role for G14-coupled proteins remains to be explored. It is worth noting that, as each LMC action potential lasts over 1 second, a reduction in any mechanism that reduces AP duration can ultimately result in an increased frequency regardless of the diastolic depolarization slope.

**Figure 23.**
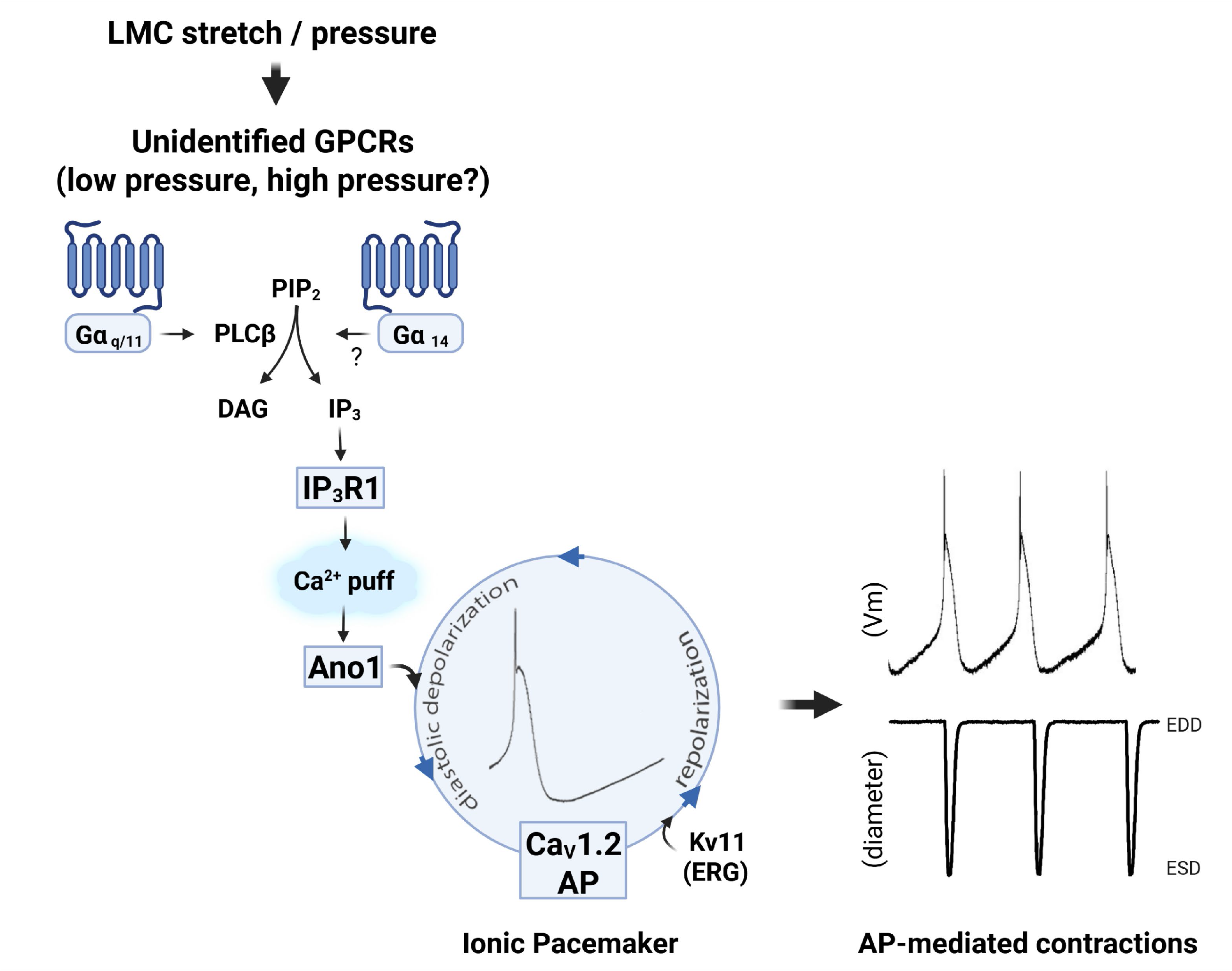
Proposed mechanotransduction mechanism underlying pressure-induced lymphatic chronotropy. Summary diagram depicting the primary signaling pathway through which changes in pressure regulate the frequency of the ionic pacemaker in LMCs. Gq/11-coupled GPCRs are the primary, but perhaps not the exclusive mechanosensor; a role for G14-coupled GPCRs, which is expressed in LMCs but not tested, is possible. Multiple GPCRs, which remain to be identified, may transduce different pressure ranges. ANO1 is the primary, but perhaps not the exclusive effector that regulates activation of Cav1.2 channels, but the pacemaker will cycle at a basal rate in the absence of ANO1 input. Kv11 provides at least one source of repolarizing signal (Kim *et al*., 2023).

### Physiological Relevance

LMC function is impaired in Cantú syndrome, the only known example of primary lymphedema produced by a gene mutation (*Kcnj8/Abbc9*) in LMCs (Davis *et al*., 2023b). This impairment is evident both in lower contraction frequencies and amplitudes at multiple pressure levels.

A lower frequency is consistent with parallel observations that gain-of-function mutations in K_ATP_ channels increase metabolic stress (Kim *et al*., 2024), whereas a lower amplitude suggests that other aspects of LMC function are altered as a consequence of chronic K_ATP_ channel hyperactivation. It is interesting that lymphatic contraction frequency and amplitude are also impaired in secondary lymphedema, in aging, and in various models of metabolic disease [(Castorena-Gonzalez *et al*., 2025) and see discussion and references in (Davis *et al*., 2023f)]. Potential therapies for reversing LMC dysfunction under these conditions will depend on identifying the key ion channels and signaling pathways that drive pacemaking and control contraction amplitude. The present study represents a major step forward in the identification of specific LMC targets that could be used for that purpose.

## Materials and Methods

### Protocol approval

All procedures were reviewed and approved by the animal care committee at the University of Missouri (protocol #9797) and complied with the standards stated in the “Guide for the Care and Use of Laboratory Animals” (National Institutes of Health, revised 2011).

### Mice

*WT* mice [C57Bl/6 (#000664) and SVF129 (#101043) strains], *Slc8a1^f/f^* (#025943) mice and *Agtr1a^-/-^*(#002682) mice were purchased from The Jackson Laboratory (Bar Harbor, Maine). Myh11-CreER^T2^ mice were gifts from Stefan Offermans (Max Plank Institute, GDR). Sperm from *Gna12^-/-^;Gna13^f/f^*and *Gna11^-/-^;Gnaq^f/f^* mice were gifts from Stefan Offermanns, after which the respective mice were rederived at MMRC, Columbia, MO to generate Myh11-CreER^T2^*;Gna13^f/f^;Gna12^-/-^*mice *and* Myh11-CreER^T2^*;Gnaq^f/f^;Gna11^-/-^*mice. *Slc8a1^f/f^* mice were crossed with Myh11-CreER^T2^ mice (in the laboratory of Joshua Scallan) to generate Myh11-CreER^T2^*;Slc8a1^f/f^*mice. *Trpc6^-/-^* and *Trpc3^-/-^* mice were gifts from Lutz Birnbaumer (NIH); *Trpc6^-/-^;Trpc3^-/-^*double KO mice were generated by crossing those two strains. Myh11-CreER^T2^*;Kcnq4^f/f^ (encoding Kv7.4)* mice were a gift from Thomas Jentsch (Free Univ. of Berlin, GDR). *Piezo1^f/f^* mice were a gift from Ardem Pataputian (UCSD) and were bred to Myh11-CreER^T2^ mice to generate Myh11-CreER^T2^*; Piezo1^f/f^* mice. *Ano1^f/f^* mice, generated in the lab of Jonathan Jaggar (Leo *et al*., 2021) were bred to Myh11-CreER^T2^ mice to obtain Myh11-CreER^T2^*;Ano1^f/f^* mice. *Itpr1^f/f^* mice were obtained from Ju Chen (UCSD) and bred to Myh11-CreER^T2^ mice to generate Myh11-CreER^T2^*;Itpr1^f/f^*mice. *Pkd2^f/f^* mice were generated in the University of Maryland PKD Core by Jonathan Jaggar, as previously described (Bulley *et al*., 2018), and bred in the Jaggar laboratory to Myh11-CreER^T2^ mice to generate Myh11-CreER^T2^*;Pkd2^f/f^* mice. *Trpm4^f/f^*mice were generated in the laboratory of Scott Earley and bred to Myh11-CreER^T2^ mice to produce Myh11-CreER^T2^*;Trpm4^f/f^*mice. *Trpv4^f/f^* mice were generated in the laboratory of Timothy Domeier (University of Missouri) and bred to Myh11-CreER^T2^ mice to produce Myh11-CreER^T2^*;Trpv4^f/f^* mice. *Cacan1c^f/f^*mice (#024714) were purchased from JAX and bred to Myh11-CreER^T2^ mice to produce Myh11-CreER^T2^*;Cacan1c^f/f^* mice. *Cacna1g^-/-^ mice* on the C57BL/6J background were a gift from Hee-Sup Shin (Korea Institute of Science and Technology), through Jeffrey Molkentin (University of Cincinnati), and were rederived into the C57BL/6J background at MMRRC (Columbia, MO). *Cacna1h^-/-^* mice were obtained from JAX (#013770) and bred into the C57BL/6J background. *Cacna1g^-/-^* mice were bred with *Cacna1h^-/-^* mice to generate *Cacna1g^-/-^*;*Cacna1h^-/-^* DKO mice (To et al. 2020). *Rosa26^LSL-mTmG^*mice were purchased from JAX (#007676) and bred to Myh11-CreER^T2^ and Myh11-CreER^T2^*;Ano1^f/f^* mice to produce Myh11-CreER^T2^*;Rosa26^LSL-mTmG^ and* Myh11-CreER^T2^*;Ano1^f/f^; Rosa26^LSL-mTmG^* mice.

For genotyping, genomic DNA was extracted from tail clips using the HotSHOT method. Genotypes were determined by PCR with 2x PCR Super Master Polymerase Mix (Catalog # B46019, Bimake, Houston, TX) according to the provider’s instructions. Mice containing Myh11-CreER^T2^ were induced with five consecutive daily injections of 100 mg tamoxifen (10mg/kg i.p.; in safflower oil) and the respective KO mice generated by crossing these to mice with floxed alleles are referred to as SM-specific (smKO) mice. Those mice were studied a minimum of 2 weeks after induction. All floxed control mice were subjected to the same induction protocol. Because the strain of Myh11-CreER^T2^ used for SM-specific deletion of floxed genes is carried on the y chromosome, only male mice were used for experiments involving Myh11-CreER^T2^mice or their floxed controls. For other strains, both male and female animals were used.

### Vessel isolation, pressure myography, and data acquisition

Mice were anesthetized with ketamine/xylazine (100/10 mg/kg, i.p.) and placed face down on a heated tissue dissection/isolation pad. The saphenous vein was exposed by a proximal-to-distal incision along the skin of the calf and the popliteal afferent lymphatic vessels on each side of the vein were isolated as previously described (Scallan & Davis, 2013). For IALVs, the skin was cut from the top of the hip to the shoulder and the skin retracted and pinned. The IALV was often found adjacent to the thoracoepigastric vein. Each vessel was then pinned with short segments of 40 µm stainless steel wire onto the SYLGARD-coated surface of a dissection chamber filled with BSA-supplemented Krebs buffer at room temperature. Once secured, the surrounding adipose and connective tissue were removed by microdissection. An isolated popliteal lymphatic vessel or IALV was then transferred to a 3-mL observation chamber, cannulated, pressurized to 3 cmH_2_O using two glass micropipettes (50-60 µm outside diameter) and moved to the stage of a Zeiss inverted microscope. Lymphatic segments used in these studies typically contained a single valve. Polyethylene tubing attached to the back of each glass micropipette was connected to a two-channel computerized pressure controller (Scallan *et al*., 2013). To minimize diameter-tracking artifacts associated with longitudinal bowing at higher intraluminal pressures, input and output pressures were briefly set to 10 cmH_2_O for popliteal or 8 cmH_2_O for IALVs at the beginning of every experiment, and the vessel segment was stretched axially to remove longitudinal slack. The lymphatic vessel was then allowed to equilibrate at 37°C with inflow and outflow pressures set to 3 cmH_2_O. Constant exchange of Krebs buffer was maintained using a peristaltic pump at a rate of 0.5 mL/min. After temperature stabilized, popliteal lymphatics typically began to exhibit spontaneous contractions within 10-15 min and stabilized in a consistent contraction pattern within ∼30 minutes. Some vessels from knock out genotypes developed insignificant or weak contractions but still developed tone; these were distinguished from damaged or leaky vessels in that the latter (which were discarded) developed neither tone nor spontaneous contractions. Custom LabVIEW programs [National Instruments; Austin, TX; (Davis, 2023a,b)] acquired real-time analog data and digital video through an A-D interface (USB-6216, National Instruments) and detected the inner diameter of the vessel at 30 fps using a Basler A641fm firewire camera (Davis, 2005). Videos of the contractile activity of lymphatic vessels were recorded for further analysis, if needed.

### Assessment of ex vivo contractile function

The contractile parameters of each vessel were characterized at different levels of intraluminal pressure spanning the physiological range from 0.5 to 10 cmH_2_O (typically in successive steps as follows: 3, 2, 1, 0.5, 3, 5, 8, and 10 cmH_2_O), except IALVs which only went to 8cmH_2_O. Spontaneous contractions were recorded at each pressure in typical intervals of 2 minutes, although a period of 4-5 min was sometimes required at the lowest pressure to obtain multiple contractions. During the pressure response protocol, both the input and output pressures were maintained at equal levels so that there was no imposed pressure gradient for forward flow. At the end of every experiment, all vessels were equilibrated by perfusion with calcium-free Krebs buffer containing 3 mM EGTA for 30 minutes, after which passive diameters were obtained at each level of intraluminal pressure.

### Contractile function parameters

Once an experiment was completed, internal diameter traces and/or 30-fps brightfield videos of spontaneous contractions were analyzed using custom-written LabVIEW programs (Davis, 2023b, a) to detect end diastolic diameter (EDD), end systolic diameter (ESD), and contraction frequency (FREQ, computed on a contraction-by-contraction basis), with each parameter averaged over a 2-5 min period. These data were used to calculate the following commonly reported parameters that characterize the contractile function of lymphatic vessels:

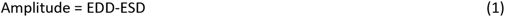

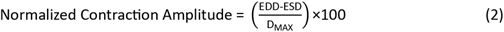

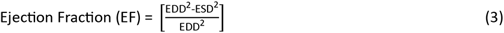

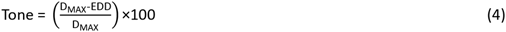

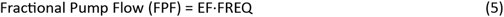

where D_MAX_ represents the maximum passive diameter (obtained after incubation with calcium-free Krebs solution) at a given level of intraluminal pressure. When frequency was zero, a value for amplitude was omitted. The relative paucity of some amplitude data at low pressures in some genotypes reflects the fact that this parameter could not be determined when frequency was zero. Each of the contractile parameters represented the average of all the recorded contractions during the measurement period at each intraluminal pressure. In popliteal vessels, data obtained for the two different periods at 3 cmH_2_O at the beginning and middle of the protocol were averaged together.

### Analysis of pressure-induced chronotropy

Quantification of the frequency-pressure (F-P) relationship for comparisons across genotypes required additional analysis steps. The average frequency for each time interval at each pressure was determined from the raw recordings of pressure and diameter using LabVIEW programs. The frequency data were then plotted as a function of pressure. Frequency tended to plateau above 5 cmH_2_O, so the subset of data in which the response was nearly linear over the pressure range 0.5-5 cmH_2_O was fitted with a first-order polynomial to obtain the slope and intercept. We considered calculating the ratio of the respective contraction frequencies at 0.5 and 5 cmH_2_O; however, in many cases no contractions occurred at P = 0.5 cmH_2_O, which would equate to a ratio of infinity. As an alternative analysis method, we computed the difference (ΔF) between the frequencies at 5 and 0.5 cmH_2_O. This approach gave consistent values that, like the F-P slope, could detect blunting of the F-P relationship, but in essentially every case the results matched those of the curve fitting procedure; thus, only the F-P slope was used for statistical analyses. To enable a more comprehensive assessment of contractile function, full plots of amplitude, frequency, FPF and tone were also plotted as a function of pressure for each genotype.

### Effects of selective GPCR inhibitors

To screen for the effects of specific GPCR antagonists, pressures were set to 3 cmH_2_O and contraction frequency was determined over a 2-5 min period. The inhibitor was added to the bath (with perfusion temporarily stopped), mixed thoroughly, and after waiting for at least 2 min, the average frequency was again determined over the subsequent 2-5 min period.

### FACS

Inguinal-axillary lymphatic vessels from tamoxifen-treated Myh11-CreER^T2^*;Rosa26^LSL-mTmG^ and* Myh11-CreER^T2^*;Ano1^f/f^;Rosa26^LSL-mTmG^*mice were dissected and cleaned of fat and connective tissue as described previously (Zawieja *et al*., 2018a). Cleaned vessel segments were transferred to a 1-ml tube of low-Ca^2+^ PSS containing (in mM): 137 NaCl, 5.0 KCl, 0.1CaCl_2_, 1.0 MgCl_2_, 10 HEPES, 10 Glucose, and 1 mg/ml BSA at room temperature for 10 min. The solution was decanted and replaced with a similar solution containing 26 U/ml papain (Sigma, St. Louis, MO) and 1 mg/ml dithioerythritol. The vessels were incubated for 30 min at 37°C with occasional agitation, then transferred to a new tube containing low-Ca^2+^ PSS with 25 mg/ml collagenase H (FALGPA U/ml, Sigma), 0.7 mg/ml collagenase F (Sigma), 20 mg/ml trypsin inhibitor (Sigma), 1mg/ml elastase (Worthington), and incubated for 6 min at 37°C. The dispersed cells were sedimented by centrifugation (300 g, 4 min), resuspended in 0.6 ml PSS containing 1mM Ca solution, and filtered through a 35-µm mesh nylon filter to obtain a single cell suspension. GFP+ LMCs were sorted by a fluorescence-activated cell sorter (FACS) using a Beckman-Coulter MoFlo XDP instrument with excitation laser (488 nm) and emission filter (530 ± 40 nm), a 70 µm nozzle, a sheath pressure of 45 psi and sort rate of 100 events per second. Sorting was performed at the Cell and Immunobiology Core facility at the University of Missouri.

### RNA isolation and quantitative, real-time PCR

Total RNA was extracted from FACS-sorted cells using the Arcturus PicoPure RNA isolation kit (ThermoFisher Scientific, Waltham, MA) with on-column DNase I treatment (Qiagen, Valencia, CA) according to manufacturer’s instructions. RNA was eluted with nuclease-free water. Purified RNA was transcribed into cDNA using a High-Capacity cDNA Reverse Transcription kit (ThermoFisher Scientific, Waltham, MA). Real-time PCR (qPCR) was performed on cDNAs prepared from each sample using 2x PrimeTime Gene Expression Master Mix (IDT, Coralville, IA) with predesigned TaqMan probes as listed in **Table 3** (IDT, Coralville, IA). Real-time PCR protocols were as follows: preheating at 95°C for 3 min, 45 cycles of two-step cycling of denaturation at 95°C for 15 sec and annealing/extension steps of 30 sec at 60°C. Data collection was carried out using a Bio-Rad CFX 96 Real-Time Detection System (software version Bio-Rad CFX Manager 3.1; Bio-Rad, Hercules, CA, USA). For analysis, the result was expressed as a ratio of target gene/reference gene (β-actin).

**Table 3.** Primers used for qPCR.

| Gene | Accession Number | Catalog Number | Description |
|---|---|---|---|
| *Ano1* | NM_178642 | IDT Mm.PT.58.12522115 | Mouse Ano1 TaqMan probe |
| *Acta2* | NM_007392 | IDT Mm.PT.58.16320644 | Mouse α-actin TaqMan probe |
| *β-Actin* | NM_007393 | IDT Mm.PT.58.33257376.gs | Mouse Actb TaqMan probe |

### scRNAseq analysis

G-protein and GPCR expression were assessed using our previously published scRNAseq dataset (Zawieja *et al*., 2025) as detailed below. This dataset was derived from a total of 10 *Rosa26^LSL-mTmG^* mice (5 males and 5 females), without Cre and without tamoxifen treatment, that were used for scRNAseq analysis. Inguinal-axillary lymphatic vessels (IALVs) from both sides of each mouse were isolated and cleaned of connective tissue, adipose tissue and associated capillaries and small arterioles. The pooled vessels were digested into a single cell suspension and the cells were kept on ice until all tissues had been processed. Cells from all vessels were combined and sorted for tdTomato expression to remove debris and concentrate the cells for creation of downstream single cell 3’ RNA-Seq libraries with 10x Genomics Chromium Chip and Chromium Next GEM Single Cell 3’ RNA-Seq reagents. Samples were sequenced with the NovaSeq 6000 S4-PE100 flow cell. Mus musculus genome GRCm39 and annotation GTF (v106) from Ensembl (https://useast.ensembl.org/Mus_musculus/Info/Index) were used to build the reference index and the reads were processed using Cell Ranger (v7.0.1) using the default parameters. The quality control and filtering steps were performed using R (v4.2.1; https://www.r-project.org/). Ambient RNA was removed from the Cell Ranger output with SoupX (Young & Behjati, 2020). A doublet score for each cell was estimated using scDBlFinder [v1.12.0; (Germain *et al*., 2021)]. Non-expressed genes (sum zero across all samples) and low-quality cells (>10% mitochondrial genes, < 500 genes, < 1,000 UMIs per cell and doublet score <0.5) were removed with custom R scripts. Cells passing filtering were normalized/scaled (SCTransformation), dimensionally reduced (t-distributed stochastic neighbor embedding (t-SNE), clustered using uniform manifold approximation and projection (UMAP), and hierarchically analyzed with Seurat (Hao *et al*., 2021; Hao *et al*., 2024) using the default parameters. Marker gene expression profiles on cell clusters and gene co-expression were visualized using Seurat and the ShinyCell R app (Ouyang et al., 2021; PMID: 33774659). The raw scRNAseq dataset is available in the NIH GEO database, #GSE277843 (Zawieja *et al*., 2025).

### Solutions and chemicals

Krebs buffer contained: 146.9 mM NaCl, 4.7 mM KCl, 2 mM CaCl_2_·2H_2_O, 1.2 mM MgSO_4_, 1.2 mM NaH_2_PO_4_·H_2_O, 3 mM NaHCO_3_, 1.5 mM Na-HEPES, and 5 mM D-glucose (pH = 7.4). An identical buffer was prepared with the addition of 0.5% bovine serum albumin (“Krebs-BSA”). During cannulation Krebs-BSA buffer was present both luminally and abluminally; however, during the experiment the abluminal solution was constantly exchanged with plain Krebs. Ca^2+^-free Krebs was Krebs with 3 mM EGTA replacing CaCl_2_·2H_2_O. All chemicals were obtained from Sigma-Aldrich (St. Louis, MO), with the following exceptions: BSA (US Biochemicals; Cleveland, OH), MgSO_4_, Na-HEPES and YM254890 (ThermoFisher Scientific; Pittsburgh, PA), ivabradine, zatebradine, S18886 (Cat. No. 5568) and losartan (Tocris). AP811, BQ123, BIBO3304 were purchased from Tocis Bioscience. YM254890, amiloride, benzamil, ivabradine, zatebradine, BaCl_2_, losartan, AP811 and BQ123 were dissolved in water to make stock solutions. BIBO3304, YM254890, Ani9, tranilast and SET2 were dissolved in DMSO to make stock solutions. Each inhibitor was then further diluted in Krebs solution to reach the final concentrations stated. Final DMSO concentrations were well below the threshold (0.4%) for direct effects.

### Statistical analysis

The data were analyzed in LabVIEW, compiled in Excel and statistical analyses were performed using Prism (v.10.2; Graphpad, San Diego, CA). IGOR (Wavemetrics, Osewego, OR) was used to read and display raw data traces. Sample sizes were determined using power analyses (Prism), with effect size for respective contraction parameters based on previously published studies by our laboratories. Statistical differences in the slopes of the F-P relationships between two groups were assessed by an unpaired, two-tailed t-test and differences between three or more groups were tested using a one-way ANOVA with Dunnett’s post-hoc tests. Comparisons between contraction parameters as a function of pressure were made using two-way or mixed-model ANOVAs with repeated measures and various post-hoc tests as indicated in the figure legends. Data are plotted as mean ± SEM. Significance levels (compared to the control group) are either stated or indicated as follows: \**p*<0.05; \*\**p*<0.01; \*\*\**p*<0.001; \*\*\*\**p*<0.0001; “n.s.” p≥0.05, with the color-coding of the symbols matching the respective group. In the figure legends, N refers to the number of animals and *n* refers to the number of vessels per group.

## Acknowledgements.

The authors are grateful for the technical assistance of Shanyu Ho and Karen Bromert. We acknowledge the kind gifts of *Kcnq4^-/-^* mice from Thomas Jentsch (Free University of Berlin), *Trpc6^-/-^* and *Trpc3^-/-^* mice from Lutz Birnbaumer (NIH), *Piezo1^f/f^* mice from Ardem Patapoutian (USCD), and *Itpr1^f/f^* mice from Ju Chen (UCSD). *Ano1*^f/f^ and Myh11-CreER^T2^*;Pkd2^f/f^* mice were gifts from Jonathan Jaggar (University of Tennessee HSC). Stefan Offermanns (Max-Planck-Institute, Bad Nauheim) kindly provided sperm from *Gna12^-/-^;Gna13^f/f^* and *Gna11^-/-^;Gnaq^f/f^* mice. The authors thank Sathish Srinivasan (OMRF) for the gift of SEW-2871. This work was supported by National Institutes of Health grants (R01-HL122578 to MJD; R01-HL168568 to JAC-G; F31HL179791 to MES; R35-HL155008 to SE; R01-HL136292 to TLD; R01-HL142905 to JPS; R01-HL143198 to SDZ).

## Data Availability

RNA sequence data were deposited in GEO (accession number GSE277843); https://www.ncbi.nlm.nih.gov/geo/query/acc.cgi?acc=GSE277843. All data generated or analyzed during this study are included in the manuscript and supporting files; source data files have been provided for all figures. LabVIEW files for data acquisition and analysis are available at the referenced Zenodo links.

---

**Figure 18-Figure Supplement 1.**
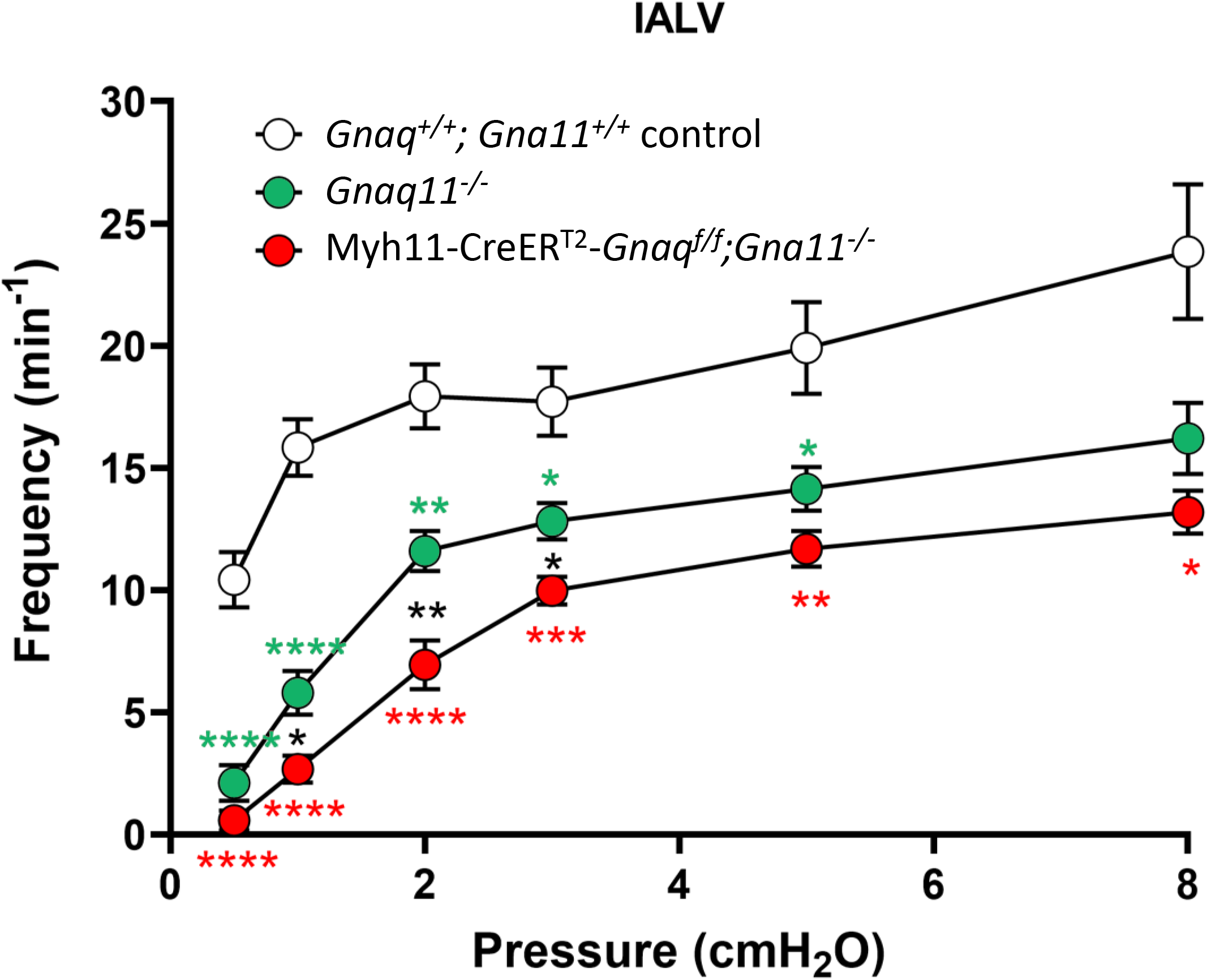
Comparisons of frequency at different pressure levels in IALVs for the various Gq/11 genotypes. Data were analyzed using a mixed model ANOVA with Fischer’s LSD post hoc tests. * = *Gnaq^+/+^;Gna11^+/+^* control vs *Gnaq11^-/-^*; * = *Gnaq^+/+^;Gna11^+/+^* control vs *Gnaq/Gna11* DKO; * = *Gna11^-/-^* vs *Gnaq/Gna11* DKO. *Gnaq^+/+^;Gna11^+/+^*control N=9, n=9; *Gnaq11^-/-^* N=13 n=13; *Gq/11* DKO N=14, n=19.

